# Segmenting and Predicting Musical Phrase Structure Exploits Neural Gain Modulation and Phase Precession

**DOI:** 10.1101/2021.07.15.452556

**Authors:** Xiangbin Teng, Pauline Larrouy-Maestri, David Poeppel

## Abstract

Music, like language, is characterized by hierarchically organized structure that unfolds over time. Music listening therefore requires not only the tracking of notes and beats but also internally constructing high-level musical structures or phrases and anticipating incoming contents. Unlike for language, mechanistic evidence for online musical segmentation and prediction at a structural level is sparse. We recorded neurophysiological data from participants listening to music in its original forms as well as in manipulated versions with locally or globally reversed harmonic structures. We discovered a low-frequency neural component that modulated the neural rhythms of beat tracking and reliably parsed musical phrases. We next identified phrasal phase precession, suggesting that listeners established structural predictions from ongoing listening experience to track phrasal boundaries. The data point to brain mechanisms that listeners use to segment continuous music at the phrasal level and to predict abstract structural features of music.

## Introduction

The physical world presents continuous streams of information, but we perceive, encode, and process it as a sequence of discrete events (Newtson et al., 1979; Zacks et al., 2007; Kurby and Zacks, 2008). The human brain segments the apparent continuity into events over multiple timescales and constructs complex hierarchical structures from natural stimuli unfolded linearly in time. This holds for spoken language, music, films, and motor sequences (Lashley, 1951; Lerdahl and Jackendoff, 1983; Ding et al., 2015; Baldassano et al., 2017). An intuitive example is that continuous speech is segmented into phonemes and syllables, which are grouped into words, phrases, and hierarchically organized sentences (Ghitza, 2012; Ding et al., 2015; Poeppel and Assaneo, 2020). Likewise, in music, units of different sizes are organized in nested hierarchical structures (Huron, 2008; Larrouy-Maestri and Pfordresher, 2018; Lerdahl and Jackendoff, 1983), such that the building blocks (i.e., notes and beats) allow motifs and phrases to be constructed according to musical criteria. Accordingly, music listening involves extracting hierarchical structures and computing long-term dependencies, comparable to spoken language processing (Maess et al., 2001; Patel, 2003; Jackendoff, 2009; Patel, 2010; Rohrmeier, 2011; Koelsch et al., 2013; Donhauser and Baillet, 2020). How abstract, high-level structure is tracked online in language is actively investigated. Evidence illuminating mechanisms of predictive event segmentation in music has, in contrast, been sparse and circumstantial. During music listening, listeners hear a single note or chord at a moment - but internally represent melodies that are stretched into the past and extended into the future (e.g. tension and attraction in musical phrases) (Augustine and Chadwick, 1991; Huron, 2008). Here we test the hypothesis that neural mechanisms well established in other domains - e.g., hippocampal physiology - underpin segmentation and prediction of high-level musical structures.

To perceive musical event structure in real time, it has been conjectured that listeners anticipate incoming musical events following past harmonic progressions as music is unfolding (Huron, 2008; Vuust et al., 2009; Rohrmeier and Koelsch, 2012; Tillmann, 2012; Patel and Morgan, 2016). Musical phrase segmentation is natural, automatic, and effortless for musicians as well as listeners without explicit knowledge of music theory (Trainor and Trehub, 1992; Kragness and Trainor, 2016). Because listening to music is ubiquitous across cultures, a deeper mechanistic understanding of this fundamental faculty is of broad significance. If abstract online musical segmentation involves fast structural predictions, neural dynamics should be observed that reflect simultaneously (i) parsing musical phrase structures and (ii) predicting phrase boundaries. Previous research has not demonstrated characteristic neural signatures of these concurrent operations.

Experiments studying online processing of music mainly focus on lower-level stimulus features, such as tracking notes or beats or rhythmicity in tone sequences (Nozaradan et al., 2011; Doelling and Poeppel, 2015; Fujioka et al., 2015; Lenc et al., 2018; Harding et al., 2019). Prediction is primarily approached on the note and beat levels, with models based on information theory or transition probability to estimate predictability of notes and chords (Pearce, 2005; Di Liberto et al., 2020). Previous studies investigating musical syntax and structural prediction typically use deviation-detection paradigms in an offline manner: neural responses to endings of well-formed musical segments (e.g. musical phrases) are compared with endings of manipulated sequences (Knösche et al., 2005; Neuhaus et al., 2006; Koelsch et al., 2013; Silva et al., 2014; Koelsch et al., 2019). Differences between neural responses to offsets of musical pieces or segments, such as the early right anterior negativity (ERAN) (Koelsch et al., 2002) or the Closure Positive Shift (CPS) (Neuhaus et al., 2006), are interpreted as evidence for processing and predicting musical syntax. It remains unclear how the brain, in real time and in natural listening settings, segments high-level musical structures and establishes fast predictions. One methodological challenge is that musical hierarchies range from a timescale of ±1 second (e.g. note, beat levels) to a timescale of ±10 seconds (musical phrase level). No neural mechanisms, especially in the time domain of tens of seconds, has been linked to processing high-level musical structures.

Here we draw on concepts from other fields. First, it has been proposed that ‘gain modulation’ in neural systems serves a fundamental role for routing and integrating information from multiple neural sources to scaffold high-level computations (Salinas and Sejnowski, 2001; Martin, 2020). In the case of music, we hypothesize that musical phrasal segmentation can be implemented through modulating the gain of low-level neural processes, such as neural tracking of notes and beats. Second, to study musical structural prediction, we draw inspiration from mechanisms of spatial cognition that have been extended to language. The phenomenon of ‘phase precession’ reveals that animals, guided by their learned cognitive maps, predict spatial positions of a future path during spatial navigation (Tolman, 1948; Jensen and Lisman, 1996; Buzsáki, 2005). Recent language research shows that prediction of high-level linguistic structures is reflected in phase precession: neural signals segmenting linguistic structures advance faster in time after listeners establish predictions for incoming speech (Teng et al., 2020). We conjecture that neural modulation components lock to musical phrasal boundaries and manifest phase precession, representing musical phrase segmentation with structural prediction.

To test these two hypotheses, we used chorales by J.S. Bach, as this music is highly structured and follows strict harmonic rules (Rohrmeier, 2011) - and collected electrophysiological data from human listeners. We first identify neural signatures that reflect the active segmentation of musical phrases. If listeners establish predictions at the phrasal level as a music piece unfolds, the neural signals of phrase segmentation should gradually peak around - and even predict - the timing of phrasal boundaries.

## Results

### Musical materials and experimental paradigm

We selected 10 Bach chorales. All music stimuli can be found at: https://osf.io/vtgse/. Nine stimuli had 8 beats per phrase, one had 12 beats per phrase. We chose three tempi, 66, 75, and 85 bpm (beats per minute) to cover a range of tempi in natural listening conditions and removed fermata that indicate any phrasal structure in the acoustics. We generated the auditory stimuli using an artificial piano sound generator, so that all beats had the same length at one tempo (see Methods). To create two distinct control conditions with disrupted phrasal structures (Fig. 1A), we (i) globally reversed the temporal order of beats in a piece by rearranging the musical scores - a ‘Global reversal’ condition, and (ii) we locally reversed the beats in the middle of musical phrases while keeping the onset and the offset beats intact - a ‘Local reversal’ condition. These reversal manipulations keep intact the basic musical contents (notes and beats) of each piece but disrupt temporally the high-level musical structures. We presented 90 pieces (10 pieces × 3 conditions × 3 tempi) to 29 participants undergoing EEG recording. The participants numerically rated how much they liked each piece (behavioral results in Fig. S1C).

**Fig. 1.**
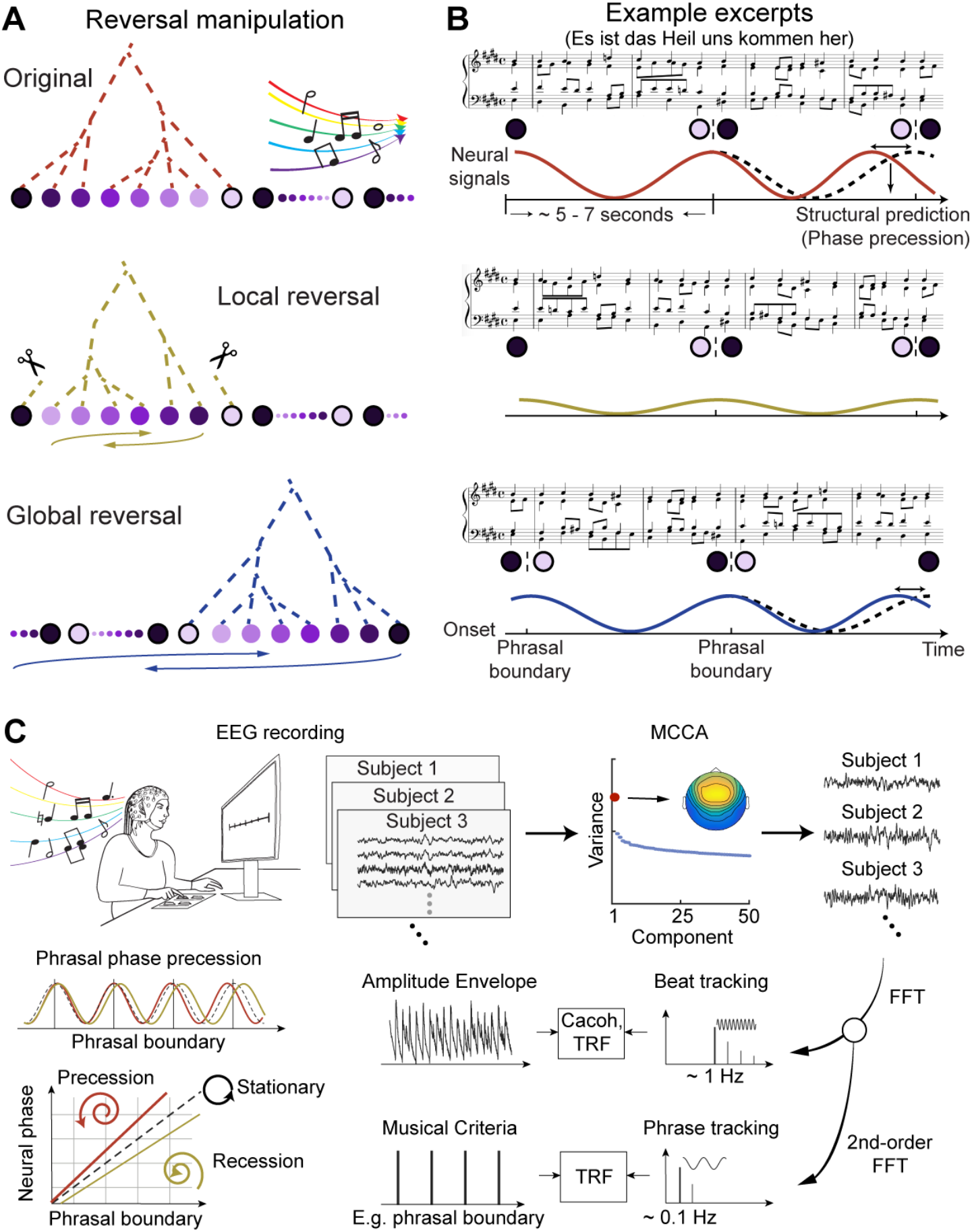
Stimulus manipulation and experimental paradigm. (*A*) Reversal manipulation. 10 Bach chorales (Original, top) were subjected to two reversal conditions: Global reversal (bottom) - the order of beats of an entire music piece was temporally reversed; and Local reversal (middle) – the middle part of the musical phrases of each piece was reversed. The harmonic progressions of musical phrases, suggested by the dashed tree structures, were hypothesized to be preserved in the Original and the Global reversal conditions but not in the Local reversal condition. (*B*) Example excerpt. The first two phrases of a piece demonstrate the reversal manipulations. Neural signals should lock to the phrasal structures in the Original (top) and Global reversal (bottom) conditions but to a lesser degree in the Local reversal (middle) condition, illustrated by the wave amplitude. Based on findings in speech segmentation (Teng et al., 2020), the preserved harmonic progressions enable listeners to anticipate phrasal boundaries at the structural level. This can be demonstrated by phrase-segmenting neural signals increasingly advancing faster than the musical structures unfolded physically – the phenomenon of phase precession. (*C*) Experimental paradigm and analysis. Participants listened to each piece while undergoing EEG recording and rated how much they liked each piece. We extracted shared neural components across the participants using multiway canonical correlation analysis (MCCA) (top right) and selected the component that explained the largest variance. We first conducted one Fourier decomposition to measure beat/note tracking around 1 Hz and derived temporal response function (TRF) and cerebral-acoustic coherence (Cacoh). We conducted the second Fourier decomposition that revealed how the power of neural signals was modulated by the phrasal structures around 0.1 Hz and derived TRFs using four different musical criteria. Lastly, we quantified phrasal phase precession in a neural-phase and phrasal-boundary plane: precession occurs when neural phase advances faster than phrasal boundaries; phase recession occurs, otherwise.

If musical phrasal structures can be extracted by listeners in the Original and Global reversal conditions (wherein harmonic progressions are preserved), neural response profiles should be observed to rhythmically lock to the phrasal boundaries, reflecting phrase-level musical segmentation. In contrast, such a phrase-segmenting neural component should be diminished in the Local reversal condition because of disrupted phrasal structures (see Fig. 1B for illustration). If listeners internally construct phrasal structures as music unfolds and establish expectations for incoming contents at the phrasal level, we should observe phrasal phase precession: neural signals locking to the phrasal boundaries advance faster than the physically unfolding music and eventually predict incoming phrasal boundaries (Fig. 1B).

### Neural tracking of notes and beats is not modulated by high-level musical structure

We first examined how the reversal manipulations modulated low-level (note and beat) music tracking (providing, as well, a replication of previous research using new stimuli). To extract the relevant neural responses, we first conducted a multi-way canonical component analysis (MCCA) over the 29 participants who generated valid EEG recordings. We derived the largest component shared across the participants (see Fig. 1C for illustration). The MCCA on the EEG signals resulted in a pooled neural signal across all EEG channels for each participant and music piece (see Methods). The weights over the EEG channels showed an auditory origin of the derived neural signals (Fig. S1E-G).

We then calculated the cerebro-acoustic coherence (Cacoh) between the amplitude envelopes of the pieces and the neural signals (Peelle et al., 2013). The results are depicted in Fig. 2A. Peak Cacoh values were observed around the beat rate (B), the note rate (N), and their harmonics, across all tempi and reversal conditions. We selected the peak Cacoh values around the beat and note rates (Fig. 2B) and conducted a three-way repeated-measure ANOVA (rmANOVA) (Condition × Tempo × Frequency, beat rate vs. note rate). We found a main effect of Frequency, with Cacoh values of the note rate significantly larger than the beat rate (*F*(1,28) = 108.09, *p* < .001, η_p_^2^ = 0.794). The main effect of Tempo was significant (*F*(2,56) = 3.68, *p* = .032, η_p_^2^ = 0.116), and the interaction effect between Tempo and Frequency was significant (*F*(2,56) = 124.47, *p* < .001, η_p_^2^ = 0.816). A two-way rmANOVA (Tempo × Frequency) showed that Cacoh increased with the tempo at the beat rate (*F*(2,56) = 26.33, *p* < .001, η_p_^2^ = 0.485) but decreased at the note rate (*F*(2,56) = 100.76, *p* < .001, η_p_^2^ = 0.783). Tempo-modulated music tracking can be explained by the resolution-integration mechanism in auditory perception (Teng et al., 2016): shortening intervals between acoustic elements facilitates global integration of acoustic information over a beat; lengthening the intervals facilitates perceptual analyses of local information to extract notes within a beat. Note that the Cacoh of note tracking is stronger than beat tracking, probably because the first harmonic of the beat note overlaps with the note rate, increasing the neural signal strength around the note rate.

**Fig. 2.**
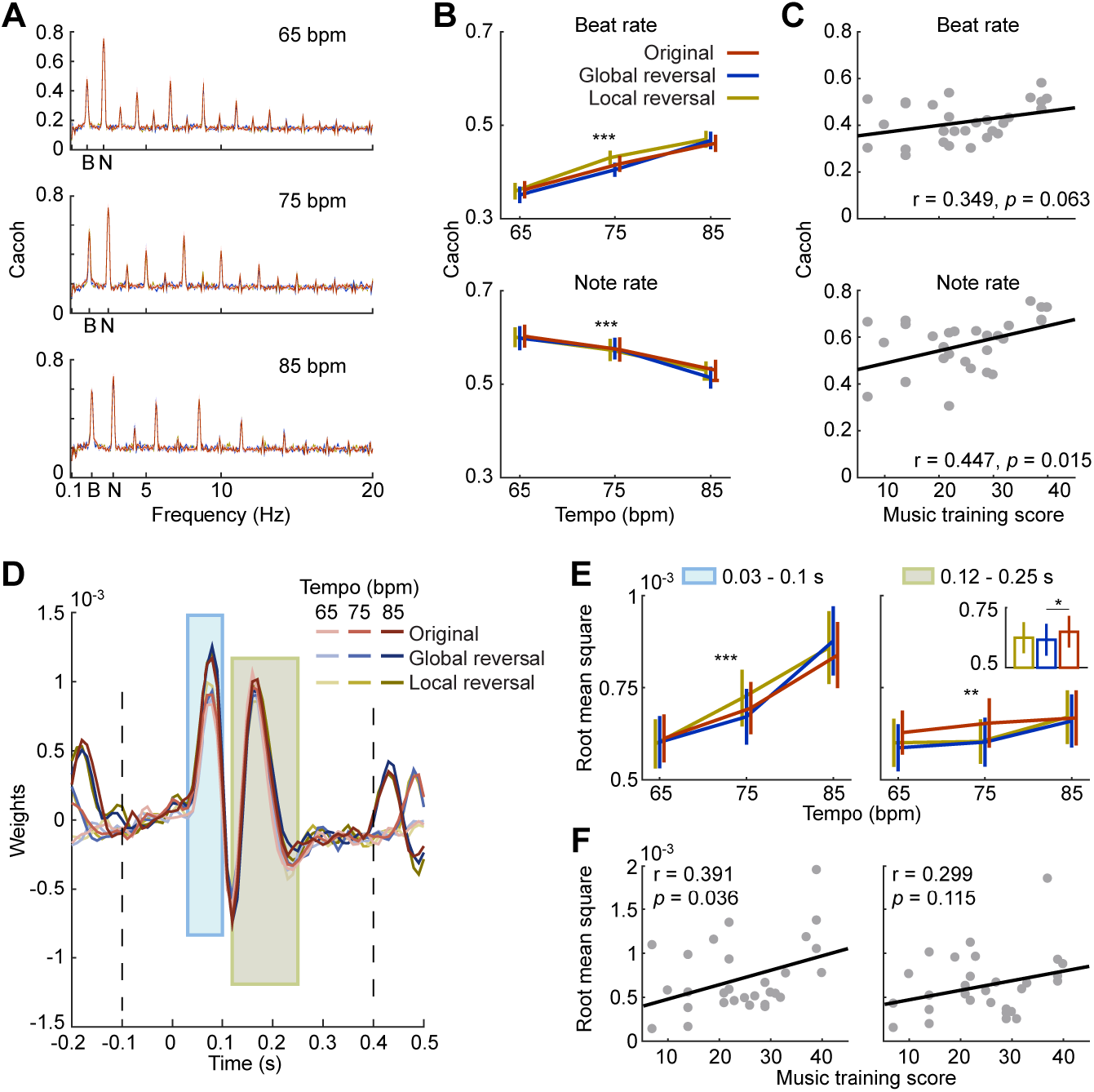
Neural tracking of beats and notes. (*A*) Cerebro-acoustic coherence (Cacoh) for notes (N) and beats (B) at each tempo. Line color indicates the reversal condition. The three conditions Original, Local, and Global are shown in each panel, but only the Original condition can be seen because the three conditions highly overlap. (*B*) Tempo modulates neural tracking of beats and notes. Increasing tempo positively modulates beat tracking (*p* < .05) but negatively modulates note tracking (*p* < .05). (*C*) Correlation between Cacoh and music training score. We used the GOLD-MSI questionnaire to quantify how much musical training each participant received and correlated the score of this subscale with Cacoh. (*D*) Temporal response function (TRF) for each condition and tempo. We identified two periods (shaded boxes) showing significant differences (*p* < .01) across all conditions and tempi. Line shade codes for the tempo. The dashed lines indicate the boundaries of the permutation test. (*E*) Root mean square (RMS) of TRF within each period. We calculated RMS within each significant period and found that the tempo positively modulated both periods (*p* < .05). In the late period, we found a main effect of conditions (*p* < .05); the RMS of the original condition is larger than the global reversal (*p* < .05). (*F*) Correlation between RMS and music training score. We correlated RMS of each period with the music training score and found a significant positive correlation in the early period (*p* < .05).

We next correlated the beat and note tracking averaged over all reversal conditions with participants’ musical training scores, obtained using the GOLD-MSI questionnaire (Müllensiefen et al., 2014) (see Methods). In line with previous research on neural tracking of musical beats and notes (Nozaradan et al., 2011; 2012; Doelling and Poeppel, 2015; Lenc et al., 2018), we show significant correlations between music training and note tracking (Koelsch et al., 2002; Fujioka et al., 2004; Doelling and Poeppel, 2015; Harding et al., 2019). We found a significant positive correlation between training and note tracking (*r*(27) = .447, *p* = .015) but not beat tracking (*r*(27) = .349, *p* = .063) (Fig. 2C). We show the correlations between the neural tracking at each tempo and the music training scores in Table S1. Surprisingly – and crucially - the main effect of Condition (Original, Local, Global) was not significant (*p* > .05) across all analyses. This suggests that previous findings of note and beat tracking may have less to do with high-level musical structures but rather reflect lower-level auditory processing of acoustic contents and tempi in musical materials.

### Reverse correlation shows temporal dynamics of music tracking

Neural tracking of notes and beats has often been investigated in the spectral domain, forgoing information on temporal dynamics (Nozaradan et al., 2011; 2012; Doelling and Poeppel, 2015; Lenc et al., 2018; Harding et al., 2019). To characterize the time domain, we derived temporal responses functions (TRF), using reverse correlation between the amplitude envelopes of the music pieces and the elicited neural signals, to uncover the dynamics in music tracking (Crosse et al., 2016). We calculated a TRF over 10 music pieces of each of nine presented versions (Fig. 2D). To reveal the modulation effects of the tempi and reversal conditions, we conducted a cluster-based rmANOVA analysis over the 9 conditions from 100 ms before the zero point to 400 ms after (see Methods). We found two significant periods (*p* < 0.01): one early period, 30 – 100 ms and one late period, 120 – 250 ms. We then calculated root mean square (RMS) representing overall signal strength over the TRF weights within each significant period, depicted in Fig. 2E. We next conducted a two-way rmANOVA (Condition × Tempo) in each period. In the early period, we found a significant main effect of Tempo (*F*(2,56) = 33.29, *p* < .001, η_p_^2^ = 0.543), with the higher tempo showing higher RMS values, but not for Condition (*F*(2,56) = .24, *p* = .788, η_p_^2^ = 0.008). This finding echoes the result of the beat tracking analysis in Fig. 2B and shows that tempo modulates early auditory responses and consequently modulates beat tracking. In the late period, we found a significant main effect of Tempo (*F*(2,56) = 6.34, *p* = .003, η_p_^2^ = 0.185) as well as, importantly, of Condition (*F*(2,56) = 4.01, *p* = .024, η_p_^2^ = 0.125). We compared the RMS of the late period between conditions and found that, after FDR correction, the original condition had higher RMS values than the global reversal (*t*(28) = 3.09, *p* = .021) (Fig. 2E, insert). We then correlated the averaged RMS across all the versions in each period with the music training score (Fig. 2F). A significant positive correlation was shown in the early period (*r*(27) = .391, *p* = .036), but not in the late period (*r*(27) = .299, *p* = .115). The correlations in all the 9 conditions can be found in Table S3.

Summarizing these first analyses, the data illustrate the basic temporal dynamics of neural responses during music listening: an early response period is primarily modulated by the tempo of music; a later period is likely related to putative musical structures. The correlations between the musical training and the neural signals are constrained to the early period (30 – 100 ms), suggesting that the observed correlation between neural tracking of beats and notes with music training in the spectral domain was mostly driven by early cortical responses, arguably related to basic auditory processing, instead of by processes underpinning musical structure tracking (Koelsch et al., 2002; 2005; Mankel and Bidelman, 2018).

### EEG power modulations at ultra-low frequencies reflect musical phrase segmentation

The effects around phrasal boundaries are likely caused by the fact that the power of the neural responses was modulated by the musical structures. By testing how the power of electrophysiological responses is modulated over the entire pieces, we sought to reveal genuine musical phrase segmentation.

We conducted a time-frequency analysis on the EEG signals to derive the spectrograms of EEG power and then calculated modulation spectra at each frequency by applying the FFT transform to the spectrograms (see Methods). We first wanted to determine whether EEG power was modulated. We show the modulation spectrum for each tempo in Fig. 3A. EEG power between 1 Hz and 2 Hz (precisely around the beat rate; y-axis) was modulated; critically, the largest modulations fell around the phrase rate (x-axis) at each tempo.

**Fig. 3.**
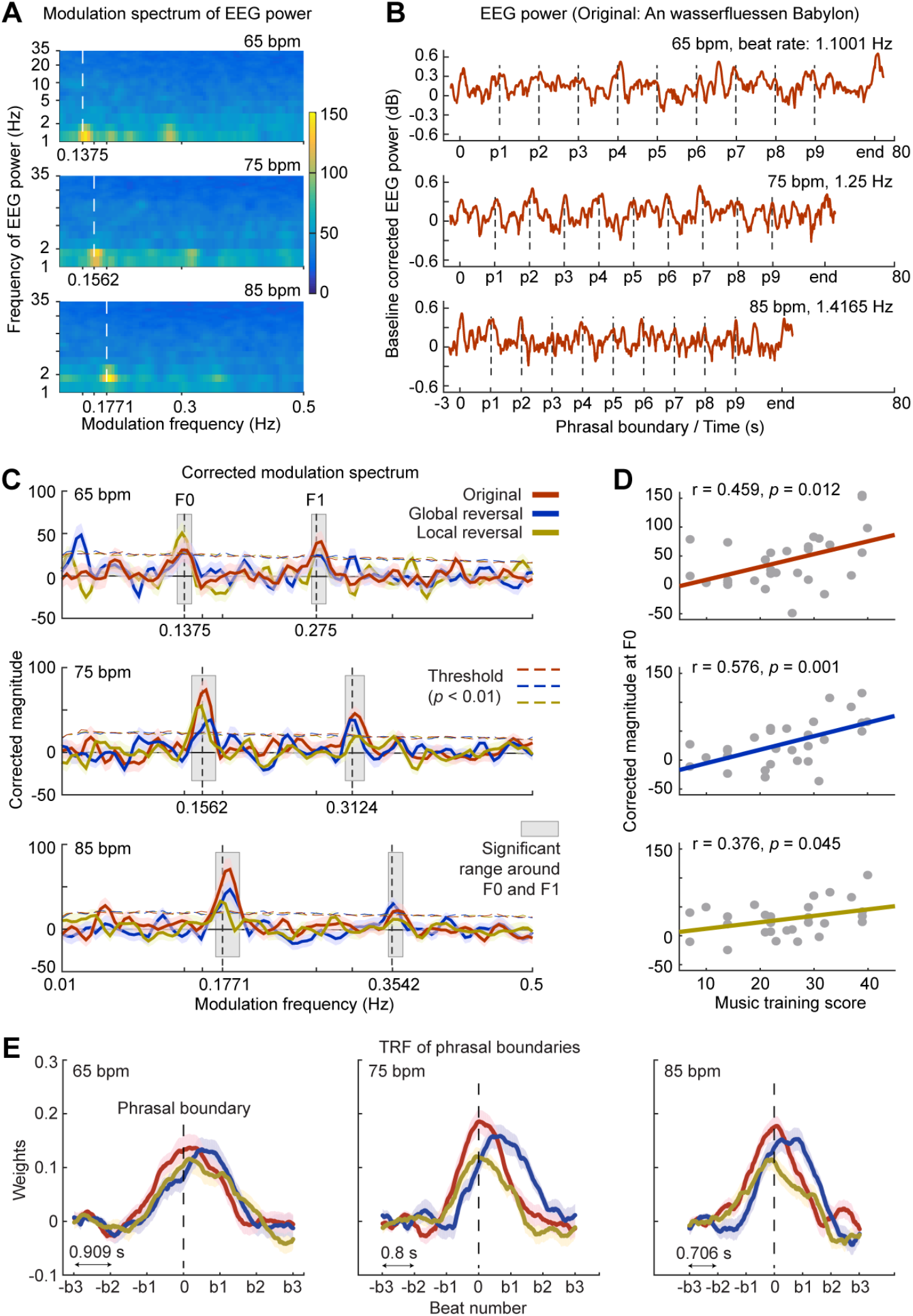
Spectral and temporal analyses of musical phrase segmentation. (*A*) Modulation spectra of EEG power. The modulation spectrum for each frequency of EEG power at each tempo was computed, averaged over the three conditions. The y-axis indicates the frequency of EEG power; x-axis the frequency of EEG power modulation. (*B*) EEG power of an example piece modulated by phrasal structure. The longest music piece in the original condition at each tempo was selected. The dashed vertical lines mark the phrasal boundaries. The fluctuations of EEG power are locked to the phrase boundaries. (*C*) Modulation spectra at the beat rate. We show the modulation spectrum at the beat rate for each condition (color code as in Fig. 1) and each tempo. The horizontal dashed lines indicate the threshold derived from the surrogate tests in the spectral domain. The shaded areas of color represent ±1 standard error of the mean over participants. The gray shaded areas represent frequency ranges with modulation amplitude above the thresholds around the fundamental frequency (F0) and the first harmonic (F1) of the phrase rate. (*D*) Correlation between musical training score and EEG power modulation of each reversal condition around F0. (*E*) TRF of phrasal boundaries. We calculated the TRF of EEG power using the phrasal boundaries as the regressor. The x axis is marked in the number of beats; the double-arrow line indicates the beat length at each tempo. Importantly, the TRFs of phrasal boundaries started to progress two beats *before* the phrasal boundaries, indicating a prediction of the phrasal boundaries. The shaded areas of color represent ±1 standard error of the mean over participants.

We extracted EEG power at the beat rate for each tempo and show the EEG power of different tempi, averaged over 29 participants, for the longest original piece in Fig. 3B. EEG Power is indeed quasi-periodically modulated by the phrasal structure, which can be seen clearly from peaks of EEG power locked to the phrasal boundaries. We next quantified the modulation spectrum of each condition for the 9 music pieces having 8 beats per musical phrase. As the lengths of the pieces were different between the three tempi, the modulation spectra cannot be directly compared across tempi, because differences of signal lengths bias estimation of their modulation spectra. Also, a baseline was needed so that we could decide the frequencies at which significant modulations of EEG power were shown. Therefore, we conducted a surrogate test in the spectral domain by shuffling the phase of each modulation frequency for each of nine reversal versions (see Methods).

Remarkably, the power modulations of the three reversal conditions were found to be significant around the corresponding phrase rate, but the strength of the modulations differed across conditions (Fig. 3C; Fig. S2A). We corrected the power modulations by subtracting the mean of the null distribution of each condition derived from the surrogate tests and then identified the significant frequency ranges around the phrase rates (F0) and their first harmonic (F1). We conducted a two-way rmANOVA (Tempo × Condition) on the corrected modulation amplitude within the significant range around the phrase rate (the fundamental frequency of the phrase segmentation, F0) (Fig. S2A). We did not find any significant main effect nor interactions (*p* > .05). We then averaged the magnitude over the significant frequency ranges around F0 of the phrase segmentation and its first harmonic (F1). The two-way rmANOVA showed a significant main effect of Condition (*F*(2,56) = 3.66, *p* = .032, η_p_^2^ = 0.115), but not of Tempo (*F*(2,56) = 0.66, *p* = .521, η_p_^2^ = 0.023) nor an interaction (*F*(4,112) = 1.48, *p* = .214, η_p_^2^ = 0.050). In the post-hoc test, after FDR correction, we did not find differences between the conditions (before FDR correction, the original is significantly larger than the local reversal, *t*(28) = 2.36, *p* = .026) (see Fig. S2A for more details).

We averaged the corrected modulation amplitude across tempi and correlated the corrected modulation magnitude of each condition around the phrase rate with the music training score (Fig. 3D). We found significant positive correlations for all the three conditions (original: r(27) = 0.459, *p* = .012; global reversal: r(27) = 0.576, *p* = .001; local reversal: r(27) = 0.376, *p* = .045). The correlations on the modulation amplitude around both F0 and F1 are shown in Fig. S2B. Note that the correlation between the musical training score and the magnitude of neural tracking of beats is not significant (Fig. 2C), but the power modulation of beat tracking significantly correlated with the musical training.

We conducted the same analysis procedure at the electrode level as well as in source space, to show topographies and source localization results (Fig. S3). Although the precision of EEG source localization was limited, it can be observed that the major difference of phrase segmentation between the reversal conditions was shown in the anterior part of the temporal lobes, with the right hemisphere showing more prominent differences than the left hemisphere, consistent with previous research suggesting a rightward lateralization of musical hierarchy processing under constrained experimental paradigms (Maess et al., 2001).

### TRFs of EEG power reveal different temporal dynamics of phrase segmentation

To capture the temporal dynamics of the phrase segmentation, we calculated TRFs of EEG power at the beat rate of each tempo using the phrasal boundaries as the regressor (Fig. 3E). The TRFs of all three conditions show peaks around the phrasal boundaries, but their weights (TRF magnitude) differed, converging with the analysis in the spectral domain (Fig. 3C&D). The weights of the TRFs of phrasal boundaries started to go above baseline well before the phrasal boundaries, indicating a *prediction* of the phrasal boundaries. To summarize the TRFs quantitatively, we fitted the group-averaged TRFs with Gaussian models (Fig. S2C&D) and tested the significance of the differences of RMS (one standard deviation of the Gaussian fits centered around the TRF peak) between the conditions by conducting a two-way rmANOVA (Tempo × Condition). We found a significant main effect of Condition (*F*(2,56) = 4.49, *p* = .016, η_p_^2^ = 0.138), but not of Tempo (*F*(2,56) = 0.93, *p* = .399, η_p_^2^ = 0.032) nor an interaction (*F*(4,112) = 0.68, *p* = .607, η_p_^2^ = 0.024). The post-hoc test shows that the Original condition is significantly larger than the Local reversal (*t*(28) = 2.96, *p* = .018, FDR corrected). Similar results were observed when we calculated RMS around the peaks of the Gaussian fits or within two standard deviations (Fig. S2E).

The TRFs of the Original and the Local reversal mostly peaked around the boundaries of the predefined phrasal structure. In contrast, the TRFs of the Global reversal had much longer latencies (Fig. S2D). This suggests that the reversal manipulations changed the phrasal structures of the music pieces in the Global reversal. We explored this finding by annotating different musical criteria that are expected to contribute to segmenting phrasal structures (i.e., cadence, grouping rule, and voice leading) in the Original and reversed music pieces, to investigate how each attribute explained the EEG power modulation (Fig. S4). The predefined phrasal structures still best explained the EEG power modulation, but the cadence showed comparable performance. The phrase segmentation in the Global reversal condition was further examined in Fig. S5.

Summarizing to this point, the findings of phrase segmentation in both the spectral and temporal domains (Fig. 3) demonstrate that we can capture how listeners parse musical phrases online. Since musical phrase segmentation was better in the Original condition than in the Local reversal condition, online phrase segmentation cannot solely be explained by the boundary beats of musical phrases, as the boundary beats in the Original pieces were matched with the Local reversal condition (Fig. 1A).

### Phrasal phase precession reveals predictive processes of musical phrase segmentation

To further explore the predictive nature of musical phrase segmentation, we quantified phase precession of EEG power modulations. The important phenomenon of phase precession has been shown in spatial navigation (Buzsáki, 2005) and related to prediction of further events (Jensen and Lisman, 1996; Lisman, 2005; Oasim et al., 2021). At the beginning of each musical piece, the neural responses will track the phrasal structure; as the musical structure unfolds, predictions for the coming phrasal structures are established and the neural phases advance faster. Because phase precession is difficult to show in continuous naturalistic stimuli, this is an opportunity to explore a fundamental neurobiological mechanism in the context of complex perceptual processing.

We quantified phrase-level phase precession for each musical piece, as the different pieces were different in terms of their musical structures, which potentially leads to different degrees of predictability among the individual pieces. One challenge with analyzing neural data from individual music pieces is that the EEG recording did not provide a sufficiently high signal-to-noise ratio for single-trial data, and it is problematic to derive robust neural indices from single trials of each participant. To address this problem, we treated the EEG data of each participant for one piece as one repetition/trial of this piece, assuming that the same piece induced similar neural responses across individual listeners. We first averaged EEG power of each music piece over the 29 participants and then quantified the phrasal phase precession on each piece. After we acquired a robust estimate on the group-averaged data, we also investigated phase precession at the level of individual participants.

Fig. 4A (left panel) shows the wrapped phase series of the group-averaged EEG power, for one music piece in the three reversal conditions at 75 bpm, plotting the phase values at the phrasal boundaries (Fig. 4A, right panel). The phase series at the phrasal boundaries in the Original and the Global reversal conditions is gradually accelerating, but not in the Local reversal. We grouped the neural phases of the music piece at the phrasal boundaries in the Original and the Local reversal conditions and visualize them in Fig. 4B (see Fig. S5 for the Global reversal). As the music was unfolding, the neural phases for the Original condition were increasingly moving forward; in the Local reversal condition, the neural phases were lagging. This can be interpreted as (perceptual) time warping in the brain (Fig. 4C, upper panel). By the end of the music piece, the neural signal indicating musical phrase segmentation predicts the future phrasal boundaries in the Original condition (Fig. 4C, lower panel). Hence, the peak frequency of phrase-segmenting neural signals should be higher than the phrase rate of the music piece in the Original condition (warped time) but lower than the phrase rate in the Local reversal condition, which is exactly what the data show in Fig. 4D (see Fig. S6 for other music pieces).

**Fig. 4.**
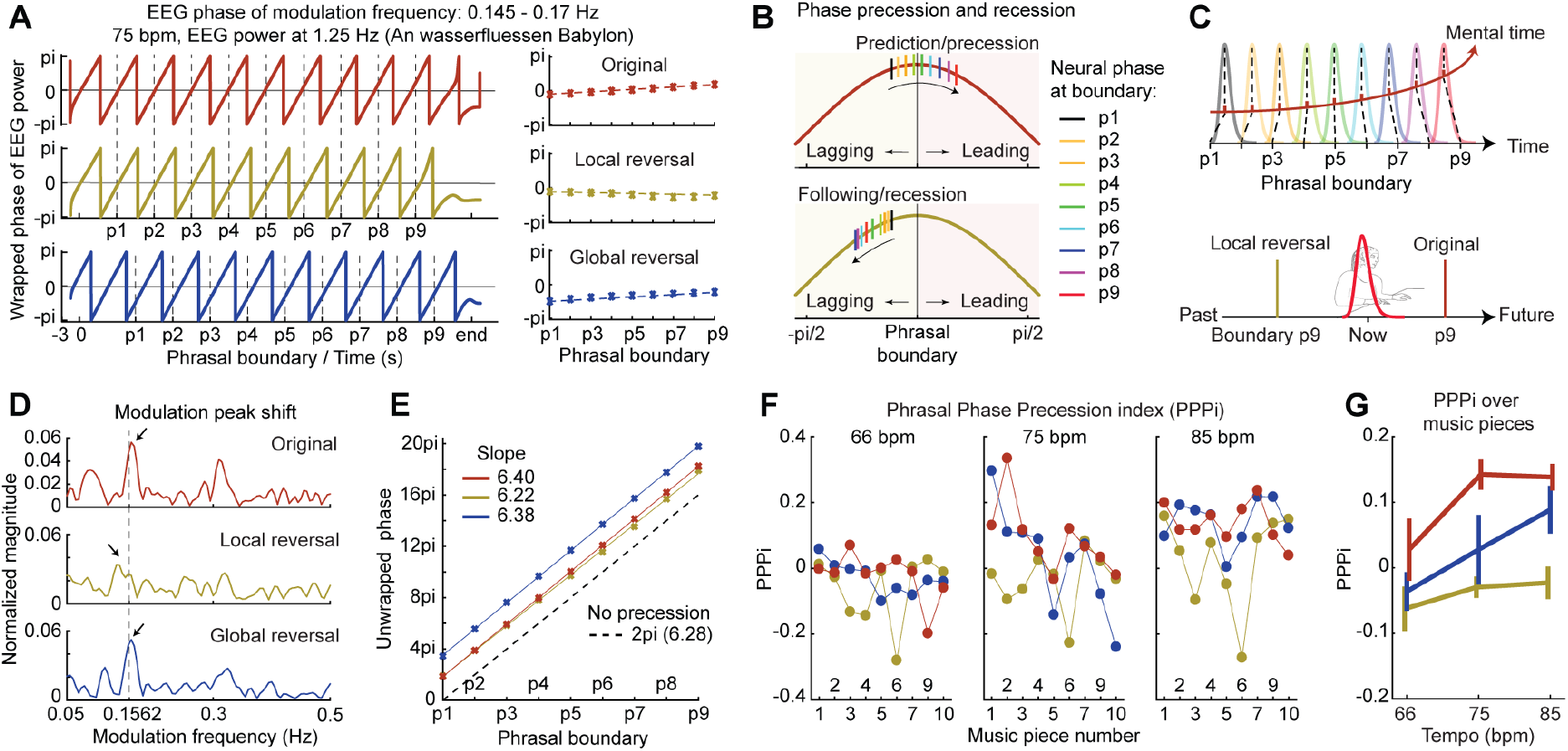
Phrasal phase precession. (*A*) Phase series of group-averaged EEG power of an example music piece (left panel). The phase of zero indicates where the peaks of cosine waves should be. The neural phases at the phrasal boundaries in the Original and the Global reversal conditions are advancing (tilting upwards) as the phrasal structures unfold (right panel). (*B*) Distribution of neural phases around phrasal boundaries. Cosine waves were plotted in both panels with the phase of zero aligning with the phrasal boundary. The bars indicate neural phases at the phrasal boundaries in the right panel of (*A*). In the Original condition, neural peaks lagged behind phrasal boundaries in the beginning but predicted phrasal boundaries after the fourth phrase; an opposite pattern was observed in the Local reversal condition. See Fig. S5 for the Global reversal. (*C*) Schematic indication of phase precession. As the phrasal phase precession occurs, time is warped mentally (top panel). Phrase-segmenting neural components followed the phrasal boundaries in the Local reversal condition but predicted the incoming phrasal boundaries in the Original condition by the end of the music piece (bottom panel). (*D*) Shifted modulation spectral peak. The peak frequencies of neural signals in the Original and Global reversal conditions are higher than the phrase rate. (*E*) Phrase Phasal Precession index (PPPi). We fit a line between the phrasal boundary number and the unwrapped boundary phase series. If no phase precession occurs, the slope is 2×pi (dashed line). The difference between the slope of each condition and 2×pi is indicated as PPPi and represents the degree and the direction of phase precession. (*F*) PPPi for each music piece. We calculated PPPi within the significant frequency ranges determined in Fig. 3C. (*G*) Averaged PPPi for each condition. The reversal manipulation modulated the phase precession (see main text). The error bars represented ±1 standard error of the mean over 9 music pieces.

We then constructed an index to summarize the phrasal phase precession by fitting a line between the order number of phrase boundaries and the unwrapped phase values for the example music piece (Fig. 4E). If no phase precession occurs, the slope of the fitted line would be 2×pi (a full cycle of neural signal for each phrase). A slope larger than 2×pi would be observed if the phase series accelerates, and vice versa. We subtracted 2×pi from the slope fitted for each piece, and the index was termed the Phrasal Phase Precession index (PPPi). Indeed, the PPPi values showed positive values for the Original version and the Global reversal of the music piece, but a negative value for the Local reversal. We quantified the effect of Tempo and reversal Condition on PPPi (shown for each piece in Fig. 4F) by conducting a two-way rmANOVA (Tempo × Condition). Each ‘participant’ here is a music piece and the rmANOVA only included 9 ‘participants’. We found a significant main effect of Tempo (*F*(2,16) = 21.65, *p* < .001, η_p_^2^ = 0.730), but no significant effect was shown for Condition (*F*(2,16) = 3.44, *p* = .057, η_p_^2^ = 0.301) nor an interaction (*F*(4,32) = 1.44, *p* = .243, η_p_^2^ = 0.153). Note that after removing the piece number 10, we found a significant main effect for Tempo (*F*(2,14) = 26.21, *p* < .001, η_p_^2^ = 0.789) and Condition (*F*(2,14) = 4.78, *p* = .026, η_p_^2^ = 0.406), but the interaction remained non-significant (*F*(4,32) = 1.33, *p* = .284, η_p_^2^ = 0.160). The above analysis on group-averaged power validated our method of using the PPPi to quantify the phrasal phase precession. We then calculated PPPi on the single trial data of each music piece at the level of individual participants (Fig. S7) and observed a significant main effect of Condition on PPPi, which was consistent with the result at the group-averaged level.

In summary, phrasal phase precession demonstrates that listeners establish structural predictions at the phrasal level from the preserved harmonic structure in the Original and Global reversal conditions during music listening and predict future event (phrasal) boundaries. The phrase segmentation in the Local reversal condition can be viewed as a passive process of phrase segmentation, as the phrasal boundaries drive neural signals, but predictive processes are only involved in the phrase segmentation in the Original and Global reversal conditions – a manifestation of musical expectation at the structural level.

## Discussion

One of the remarkable features of perceptual and cognitive systems is that we can track not just surface properties of naturalistic stimuli but also extract higher-order structural features which are not obviously visible. This capacity is especially prominent when we process continuously varying, complex stimuli such as speech and music. Investigating these internal, inferential processes has proved to be challenging. We discovered a neural signature that segments musical phrases online, in a manner that exploits predictive processes demonstrated by phase precession.

We first analyzed neural tracking of notes and beats in the spectral domain (applying novel approaches to a well-studied question). Importantly, we found that the note and beat tracking did not interact with the presence or integrity of phrasal musical structures (Fig. 2A, B&C). The subsequent temporal analysis showed that the neural tracking had two characteristic periods: an early period modulated by tempo and a late period sensitive to phrasal structure in music (Fig. 2D, E&F). We then demonstrated that EEG power of the beat tracking was modulated by musical structure and the power modulation reflected *musical phrase segmentation* (Fig. 3). The analyses culminate in the quantification of phrasal phase precession. We show that musical phrase segmentation is a predictive process rather than being passively evoked by the phrasal boundaries (Fig. 4). Our findings support the view that listeners actively segment music streams online, conforming to high-level musical structures.

The approach show that it is now possible to move beyond the neural tracking of notes and beats, lower-level features in music, and study fundamental (and relatively abstract) musical structures realized at long timescales (> 5 s), using newer analysis approaches to non-invasive EEG recording. Previous studies have typically shied away from analyzing EEG signals at ultra-low frequencies (i.e. ∼ 0.1 Hz) due to the low signal-to-noise ratio. However, the neural signatures of phrase segmentation we observed were reliable and could be estimated from a single music piece, allowing the possibility to study differences between individual music pieces (Fig. 3B and Fig. 4A&F). Although we were motivated by previous research showing characteristic responses to the endings of manipulated musical structures, such as ERAN (Koelsch et al., 2002) and CPS (Neuhaus et al., 2006), our new approach reveals neurobiological insights about music listening in natural conditions as well as neural signatures of phrase segmentation that differ in kind from ERAN and CPS. (1) The neural signature of processing the musical syntax can be directly observed (Fig. 3B&C), instead of by comparing different conditions of musical materials, as in ERAN and CPS. (2) The processing of the phrasal structures started not after the ending of a musical segment but before the phrasal endings (Fig. 3E). (3) The phrase segmentation derived from the low-level neural tracking of beat, modulated by high-level musical structures. More importantly and generally, we show that segmenting sensory (music) streams based on high-level grouping structures is a dynamic operation that exploits predictive processes (Fig. 4).

The phenomenon of phase precession has been argued to reflect predictions of future events (Jensen and Lisman, 1996; Lisman, 2005). The *phrasal phase precession* we show – the discovery that neural phase accelerates as music unfolds – illustrates that the structure-depended segmentation of music streams is not solely based on the musical stimuli but driven by listeners’ active construction of a segmentation schema (Newtson et al., 1979). Listeners potentially extract structural regularities in music from the beginning of a musical piece and gradually internalize the pattern of structure of the musical pieces, which allows listeners to construct a template to segment the musical piece. Hence, listeners can better segment the phrasal boundaries as the music is unfolding. This is very much analogous to the phenomenon of phase precession in the spatial domain – after being familiarized with mazes, animals rely on memorized spatial landmarks to predict the path ahead, which results in phase precession of neural signals (Buzsáki, 2005). We here provide the evidence that segmentation is an interplay between stimulus-driven processes and active mental construction of future events.

The musical phrase segmentation we report probably reveals a neural signature of segmenting high-level musical structures in general. The original music pieces we used were selected so that they had clear and simple musical phrasal structures, coherent across different music theoretical criteria, such as voice leading, group-changing rule, and cadence (Fig. S4B&C). This selection facilitated the successful identification of the neural signature of musical phrase segmentation, but the exact contribution of each criterion should be specifically investigated. In any case, the shifted latency of TRF of the power in the Global reversal condition (Fig. 3E&S5) indicates that phrase segmentation varied with the music criteria, modified by the reversing procedure. Next steps should include disentangling the neural modulations of different musical criteria to reveal the precise computational nature of the musical segmentation of high-level structures we identified (Fig. S4 and Fig. S5).

Musical phrase segmentation resides in a timescale of above ∼5 seconds (the minimum phrase length at 85 bpm), requiring recruitment of brain areas capable of processing information over a long timescale. Although the EEG signals we extracted using MCCA mostly related to neural activities of an auditory origin (Fig. S1D&F), the auditory system does not typically represent such long time constants (Overath et al., 2015; Teng et al., 2017; Teng and Poeppel, 2019; Donhauser and Baillet, 2020) and hence is not equipped to extract long-term phrasal structures. The candidate brain regions with long time constants are frontal areas, which have been proposed to integrate information over a long time period in language processing (Hagoort et al., 2004; Hagoort, 2005; Hickok and Poeppel, 2007; Lerner et al., 2011; Hagoort and Indefrey, 2014) and movie processing (Hasson et al., 2008), and have been demonstrated to be involved with processing musical syntax (Maess et al., 2001; Knösche et al., 2005; Koelsch et al., 2005). It is likely that the frontal areas (e.g. potentially right inferior frontal areas as suggested in Fig. S4B) establish the phrasal structures online and convey signals to modulate the neural responses of beat tracking. On the other hand, with regard to predicting phrasal event boundaries (Fig. 4), the hippocampus likely aids in encoding past musical structural information and fast retrieval of structural knowledge before phrasal boundaries for prediction (Baldassano et al., 2017; Ben-Yakov and Henson, 2018). Lastly, predicting the timing of auditory sequences has been shown to involve motor areas, including premotor cortex, supplementary motor area, and the basal ganglia (Grahn and Rowe, 2012; Morillon and Baillet, 2017; Assaneo and Poeppel, 2018; Rimmele et al., 2018; Assaneo et al., 2019). Hence, we conjecture that a fast interplay between frontal areas and hippocampus during music listening supports extracting event structures and establishing structural predictions online, with assistance from motor areas for predicting the precise timing of musical events. It would be interesting to test these hypotheses by employing MEG and/or functional MRI to pinpoint the brain areas involved in the musical phrase segmentation for a more comprehensive understanding.

We identified a robust neural signature that captures the online segmentation of music according to its high-level phrasal structure. The novel quantification of phrasal phase precession further demonstrates a predictive process of phrase segmentation during music listening. The neural signatures we highlight (musical phrase segmentation and phrasal phase precession) and the analysis procedures (EEG power modulation and PPPi) provide novel directions for studying cognitive processes of high-level structures using non-invasive recording techniques such as EEG or MEG.

## Methods

### LEAD CONTACT AND MATERIALS AVAILABILITY

Further information and requests for resources should be directed to and will be fulfilled by the Lead Contact, Xiangbin Teng.

#### Participants

Thirty-four native German speakers (age 18 – 34, 23 females) took part in the experiment. All the participants self-reported normal hearing and no neurological deficits. For the results of liking rating on musical materials, we included 32 participants (age 18 – 34, 21 females), because two participants could not finish the experiment (one participant chose to terminate the experiment, and we encountered technical issues when testing the other participant). For the EEG recording, 29 participants were included (age 18 – 34, 20 females). Among the five participants excluded from EEG analyses: one participant did not undergo EEG recording; we encountered technical issues during EEG recording for two participants; and two participants could not finish the experiment. Written informed consent was obtained from each participant before the experiment and monetary compensation was provided after the experiment. The experimental protocol was approved by the Ethics Council of the Max Planck Society.

#### Individual scores of musical training

In order to investigate the relation between the musical training of the participants and their neurophysiological responses to music listening, we used the Goldsmiths Musical Sophistication Index (GOLD-MSI) (Müllensiefen et al., 2014). This self-report inventory (German version) was given to the participants before the EEG recording and includes six dimensions: Active Music Engagement, Self-reported Perceptual Abilities, Musical Training, Self-reported Singing Abilities, and Sophisticated Emotional Engagement with Music. We mainly focused on Musical Training here.

#### Stimuli

We selected 11 music pieces from the 371 four-part chorales by Johann Sebastian Bach (Breitkopf Edition, Nr. 8610). In the 18^th^ century, chorales were congregational hymns performed in religious settings. The regularity and transparency of the harmonic structures of the Bach chorales facilitated our subsequent analyses in a frequency tagging paradigm to investigate musical phrase segmentation. Musical phrases are generally marked by fermatas, indicating a pause or the prolongation of a note (Fuller, New Grove Dictionary of Music Online; Schilkret, 1988); the predefined phrasal structures of the selected pieces are shown in Table S1.

We then processed the original musical scores so that several confounds that may appear in EEG analyses could be avoided. We first ensured that each beat was always marked by a note onset, and hence the neural tracking of beat could be measured accordingly. Concretely, the whole and half notes were substituted by four or two quarter notes and the ties across notes were deleted that they could be repeated. The material consisted thus mainly of quarter and eighth notes and few sixteenth notes (which were allowed because they occurred at regular positions within and between music pieces). More importantly, as fermatas in the original music pieces provided rhythmical or acoustic cues for phrasal structures (such as pauses between phrases and lengthened notes), we removed them so that listeners could only rely on harmonic structures as well as other structural cues, such as cadence and voice leading, to parse music streams into phrases. The processed musical scores can be found in a public data repository (https://osf.io/vtgse/).

In order to investigate musical phrase segmentation, it is necessary to provide two control conditions: (1) phrasal structures in each music piece are disrupted but basic musical contents, such as notes and tempi of beats, are kept intact; (2) phrasal structures are preserved to a large extent but listeners’ familiarity with music materials is controlled. For this purpose, we employed a paradigm of temporal reversal that has been commonly used in speech and language studies (Saberi and Perrott, 1999; Baldassano et al., 2017).

We first applied a *local reversal procedure* to the music pieces. According to the phrasal structures defined in the musical scores, we kept the onset beat and the ending beat of each phrase intact but temporally reversed the order of the beats in the middle. For example, a phrase of eight beats [1 2 3 4 5 6 7 8] becomes a new phrase [1 7 6 5 4 3 2 8] after the local reversal. A schematic illustration of this reversing procedure on the musical scores can be seen in Fig. 1A. The rationale for this local reversal was as follows: as we kept the salient onset and ending beats intact but disrupted the phrasal structures, neural indices for tracking musical phrases should be lower in the condition of local reversal than in the original condition. Alternatively, if the salient onset and ending beats are sufficient for listeners to parse continuous music streams into musical phrases, musical phrase segmentation should not differ between such locally reversed music pieces and the original pieces. To implement this local reversal procedure, we first used MuseScore 3 (musescore.org) to generate audio files of the music pieces from the processed original scores. As the beats in the generated audio files had the exact same length at a certain tempo, we cut out each beat and then reversed the order of the beats while preserving the waveform of each beat (the note order in a beat was not reversed).

Secondly, we applied a *global reversal procedure* on the music pieces. For each piece, the order of all the beats was temporally reversed, starting from the end to the beginning. A schematic illustration of this procedure can be seen in Fig. 1A. This reversing procedure was implemented on the order of beats but not on the waveforms of beats. By reversing the music pieces globally, we aimed to preserve harmonic progressions and the phrasal structures but to disrupt the typical onset and end beats of musical phrases. If musical phrase segmentation can still be observed for the globally reversed pieces, and if the neurophysiological tracking of musical phrases is comparable to the original pieces, it can be concluded that harmonic cues play an essential role in neurophysiological tracking of musical phrases. Also, as globally reversed music pieces are not typically heard by listeners, the listeners’ familiarity of the music pieces can be shown to be a minor contributor to musical phrase segmentation, if the musical phrase segmentation can still be observed.

Note that the *harmonic progression* was particularly disrupted in the Local reversal condition whereas the Global reversal manipulation allowed to keep a plausible articulation of chords over time (Fig. 1A). We extracted the amplitude envelopes of the pieces and showed the modulation spectra of the envelopes in Fig. S1A. The acoustic differences between the three reversal conditions (Original, Local reversal, Global reversal) around the phrase rates are negligible (Fig. S1B), so we conjecture that the differences of neural signals between the reversal conditions must be due to differences in high-level musical structure.

We generated the music audio files in MuseScore 3 for three tempi, 66 bpm (beats per minute), 75 bpm, and 85 bpm, which arguably cover a reasonable range of tempi of actual recordings of the Bach chorales (the tempo range we observed in a set of forty-five interpretations by voice, organ, or piano). Choosing three tempi, instead of only one, also enabled us to investigate how musical phrase segmentation varied with tempo. Furthermore, if phrase and beat tracking are observed in the music pieces of different tempi at different frequencies of EEG signals, this aids in validating the neural measurement – neural tracking of music structures is not constrained to a specific tempo or a frequency range of EEG signals but aligns with musical structures.

In summary, we had three reversal conditions - original (without reversal), global reversal, and local reversal, and three tempi - 66 bpm, 75 bpm, and 85 bpm. This generated a 3-by-3 experimental design. We used the music piece number 0, BWV 255 (Table S1), as the training piece before the formal experiment to familiarize participants with the experimental materials and the task. The 10 remaining pieces were included in the data analyses. In total, we presented 90 music pieces (3 conditions * 3 tempi * 10 pieces). The sampling rate of audio files was 44,100 Hz and the amplitude of the audio files was normalized to 70 dB SPL by referring the music materials to a one-minute white noise piece that had the same sampling rate and was measured beforehand to be 70 dB SPL at the experimental setting. The music stimuli used in the current study can be found in a public data repository (https://osf.io/vtgse/).

#### Analysis on acoustics of the music materials

To characterize temporal dynamics of the music pieces, we derived the amplitude envelopes of the musical materials. We filtered the audio files through a Gammatone filterbank of 64 bands logarithmically spanning from 50 Hz to 4000 Hz (Ellis, 2009). The amplitude envelope of each cochlear band was extracted by applying the Hilbert transformation on each band and taking the absolute values (Glasberg and Moore, 1990). The amplitude envelopes of 64 bands were then averaged, and we downsampled the sampling rate from 44100 Hz to 100 Hz to match the sampling rate of the following processed EEG signals for further analyses (i.e., cerebral-acoustic coherence and temporal response function; see below).

We transformed the amplitude envelopes to modulation spectra so that the canonical temporal dynamics of the music pieces can be concisely shown in the spectral domain. We first calculated the modulation spectrum of the amplitude envelope of each music piece using FFT with zeropadding of 8000 points and took the absolute value of each frequency point. As each piece had a different length and the zero-padding caused the modulation spectra of different music pieces to have different ranges of magnitude, we normalized the modulation spectrum of each piece by dividing the norm of its raw modulation spectrum. We then averaged the normalized modulation spectra across the 10 pieces for each condition at each tempo. At each tempo, the average modulation spectra across three conditions were highly similar, so we only showed the modulation spectra averaged across the three conditions for each tempo (Fig. S1A). Nonetheless, to demonstrate that the differences of the amplitude modulation spectra between the three conditions were not noteworthy, we calculated at each tempo the standard deviation over the modulation spectra of three conditions (Fig. S1B). The music pieces of the three conditions showed the standard deviations close to zero below the beat rate, so the neural tracking of phrasal structures found later cannot be caused by the acoustic differences between the music stimuli of different reversal conditions.

#### Analysis of music structure along four criteria

Although it can be assumed that the phrasal structure of the music pieces corresponds to the structure illustrated by the fermatas, it is likely that different musical criteria contribute to segmenting the music pieces and thus support musical phrase segmentation. To investigate how listeners parse music streams using the predefined phrasal structures as well as different musical criteria, we again borrowed a methodology used in the studies on speech and language wherein linguistic boundaries of various levels in the speech or language materials are marked according to linguistic theories and then are used (e.g. as regressors) to study the neural signatures around such linguistic boundaries (Di Liberto et al., 2015; Brodbeck et al., 2018; Broderick et al., 2018). Two musicians with training in music theory manually marked musical structures of the musical materials we generated according to three criteria: voice leading (Trainor et al., 2014), cadence, and grouping rule (Deliege, 1987). The two musicians agreed on their decisions on the markings of the musical structures. We show the musical structures determined by voice leading, cadence, and grouping preference rule in Table S2.

#### Experimental protocol and EEG recording

EEG data were recorded using an actiCAP 64-channel, active electrode set (10–20 system, Brain Vision Recorder, Brain Products, brainproducts.com), at a sampling rate of 500 Hz, with a 0.1 Hz online highpass filter (12 dB/octave roll-off). There were 62 scalp electrodes, one electrode (originally, Oz) was placed on the tip of the nose. All impedances were kept below 5 kOhm, except for the nose electrode, which was kept below ∼10 kΩ. The auditory stimuli were delivered through plastic air tubes connected to foam earpieces (E-A-R Tone Gold 3A Insert earphones, Aearo Technologies Auditory Systems).

The experiment included a training session and a testing session. In the training session, we presented the three conditions of the piece BWV 255, at the three tempi, to the participants, who were instructed to answer a question after listening to each piece: ‘how do you like this music piece’. The participants rated the music pieces using a 6-point scale from 1 to 6, with 6 being the most positive rating and 1 being the most negative rating. In each trial, before each piece was presented, the participants were required to focus on a white cross in the center of a black screen. After 3 to 3.5 seconds of silence, a piece was presented and the participants were instructed to keep their eyes open while listening to it. After each piece ended, the question appeared on the screen and the participants rated the piece. The next trial started right after the participants’ responses. The order of music pieces was randomized between participants. The behavioral data and EEG signals were not recorded during the training session.

The testing session followed the training session. We presented the 90 music pieces in six blocks to the participants while they were undergoing EEG recording. 15 pieces were presented in each block. The order of the music pieces was randomized within and across the blocks, and a different order of pieces was presented to each participant. The trial structure was the same as in the training session. Behavioral ratings were recorded. After each block, the participants could choose to take a short break of around one to three minutes or to initiate the next block.

#### Behavioral data analysis

Behavioral data were analyzed using MATLAB 2016b. To quantify differences between conditions and tempi, at each tempo for each condition for each participant, we averaged liking ratings over 10 music pieces and used the mean as the measurement for each tempo and each condition on which the statistics were conducted. The results indicated how each condition and each tempo affected the liking ratings.

#### EEG preprocessing

EEG data analysis was conducted in MATLAB 2016b using the Fieldtrip toolbox 20181024 (Oostenveld et al., 2011), the wavelet toolbox in MATLAB, NoiseTools (de Cheveigné et al., 2018; 2019), and the Multivariate Temporal Response Function (mTRF) Toolbox (Crosse et al., 2016).

EEG recordings were off-line referenced to the average of activity at all electrodes. Raw EEG data were first filtered through a band-pass filter from 0.5 to 35 Hz embedded in the Fieldtrip toolbox (a FIR zero-phase forward and backward filter using MATLAB ‘fri1’ function with an order of 4). Each trial (the recording for each music piece) was divided into an epoch of a length of each music piece plus a 3 second pre-stimulus period and a 3 second post-stimulus period. Baseline was corrected for each trial by subtracting out the mean of each trial, which was necessary for the following multiway canonical correlation analysis (MCCA) (de Cheveigné et al., 2018; 2019). An independent component analysis was applied for EEG recording of all the trials and used to correct for artifacts caused by eye blinks and eye movements.

#### Multiway canonical correlation analysis (MCCA)

We focused on the neurophysiological signals evoked by the musical pieces (auditory stimuli). To extract EEG signals that largely reflect auditory-related neural responses, instead of arbitrarily selecting certain electrodes (*e.g.* CZ or FCZ), we deployed MCCA to extract shared EEG components across all the participants. The participants were listening to the same set of pieces and hence similar neural responses to the music pieces should be observed across participants, although each participant’s EEG recording varied with his or her head size and EEG cap position and with different sources of noise. MCCA, briefly, first implements a principal component analysis (PCA) on each participant’s EEG recordings and then pools all the PCA components across the participants to conduct another set of PCA. The components from the second PCA reflect shared components of neural responses across the participants that are invariant to EEG cap position and individual head sizes. Detailed procedures and further explanations can be found in de Cheveigné et al. (2019). This procedure of component extraction simplified further analyses and avoided biases introduced by arbitrary EEG channel selections and by differences of EEG cap positions and head sizes across participants.

We first sorted each participant’ EEG recordings according to the music piece number and concatenated EEG recordings of each condition and each tempo after epoching to form one long trial. For example, the EEG recording for the piece 1 (BWV153/1) was put in the beginning and was followed by the EEG recordings of the piece number 2, 3, 4, 5, 6, 7, 8, 9, and 10, sequentially. The 3 second pre- and post-stimulus periods were included for each trial. For each condition and each tempo, we applied MCCA to derive 50 components and selected the first component, as the first component explaining the most variance (Fig. S1D&E). We checked MCCA weights over EEG channels and plotted them as topographies in Fig. S1D, from which it can be seen that the topographies of MCCA weights resembled typical topographies of auditory EEG responses. To validate our component selection, we calculated the power spectra for the first five components of MCCA in one condition (local reversal, 65 bpm). We zero-padded the time series of each component to 8000 points and calculated power spectra using FFT. We took the absolute value of each frequency point and normalized the amplitude spectrum by its norm. The results are shown in Fig. S1F&G, from which it can be seen that only the first component showed amplitude peaks corresponding to the frequency of the beat rate of each tempo. This demonstrated that the first component of MCCA indeed represented neural responses to music pieces.

After conducting MCCA for each condition and each tempo and extracting the first component, we projected the component back to each participant and derived one long trial including the 10 music pieces for each participant. We then cut out the neural response for each piece from the long trial. As PCA sometimes reversed polarity of EEG signals, the polarity of the derived signals was manually checked and corrected for each participant.

#### Cerebral-acoustic coherence (Cacoh)

We calculated the coherence between the neural signals and the amplitude envelopes of music pieces in the spectral domain - *cerebral-acoustic coherence (Cacoh)* - to investigate how pieces of different conditions were tracked in the brain at different neural frequencies. The rationale is that, if the music pieces of one condition can be more reliably tracked by the auditory system than the pieces of the other conditions because of their specific musical structures or tempi, this difference can be observed in the magnitude of coherence between the neural signals and the amplitude envelopes. This measurement, Cacoh, has been used in neurophysiological studies on speech perception (Peelle et al., 2013; Doelling et al., 2014), which quantified how phase-locked neural responses correlate with acoustic signals – at what frequencies and how strong the neural signals and the acoustic signals correlate with each other. In essence, Cacoh calculates cross spectrum coherence between neural signals and acoustic signals; here we calculated magnitude-squared coherence using the function ‘mscohere’ in MATLAB 2016b (https://de.mathworks.com/help/signal/ref/mscohere.html). This allowed us to control the temporal window used to calculate the cross-spectrum coherence and the spectral resolution of the coherence values so that the temporal window sizes and the spectral resolution were matched across all the conditions. This helped avoid the biases of calculating spectral coherence over the entire music pieces introduced by different lengths of the music pieces. Nonetheless, we still referred to our calculation as ‘Cacoh’ as this term has been well adopted in the literature.

We calculated Cacoh for each music piece and each participant using the neural data and the amplitude envelope derived from ranging from 1 second after stimulus onset to 1 second before offset, to minimize the influence of onset and offset neural responses of a whole musical piece. The temporal window used was a Hanning window of 1024 points with an overlap of 512 points (50 percent of overlap between adjacent temporal windows). We constrained the frequency range from 0.1 to 20 Hz, with a step of 0.05 Hz. The inputs, the neural signals, and the amplitude envelope had a sampling rate of 100 Hz. We then averaged Cacoh values across the 10 pieces within a condition. Therefore, a Cacoh value was derived for each tempo and each condition at each frequency point.

#### Temporal response function (TRF)

To investigate the temporal dynamics of the phase-locked responses to the music pieces, we calculated the reverse correlation between the neural signals and the music amplitude envelopes in the temporal domain – temporal response functions (TRFs). TRF has been commonly used to investigate how the auditory system codes acoustic and linguistic features in neurophysiological studies on speech perception (Di Liberto et al., 2015; Brodbeck et al., 2018; Broderick et al., 2018). The rationale for this analysis is similar to Cacoh: if music pieces of a condition robustly evoke auditory responses, such robust auditory responses should be reflected by TRF in the time domain. Furthermore, we could extract temporal information from TRFs on when the auditory responses are modulated by different conditions. For example, TRFs may reflect differences between tempi within the first 100 ms after the onset of an auditory event while TRFs may differ between the conditions after 100 ms because of different manipulations on the musical structures. Early auditory responses likely reflect sensory processes whereas late responses tend to reveal high-level cognitive processes in music perception (Koelsch et al., 2002; Koelsch and Siebel, 2005; Koelsch et al., 2005).

The TRF was derived from the amplitude envelopes of stimuli (*S*) (for details see *Analysis on acoustics of stimuli*) and their corresponding EEG signals (*R*) (for details see *EEG preprocessing and analysis*) through ridge regression with a parameter (lambda) to control for overfitting (superscript *t* indicating transpose operation):

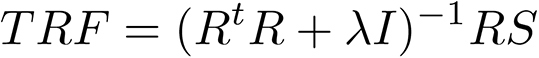

We calculated a TRF for each music piece for each participant using the neural data and the amplitude envelopes between 1 second from the stimulus onset to 1 second before the offset so that the influences of the onset and offset neural responses were avoided. The TRFs were calculated from 200 ms before onsets of auditory events and 500 ms after. As the highest tempo was 85 bpm, the shortest inter-beat interval of the music pieces was around 700 ms and the inter-note interval was less than 400 ms. In the statistical analysis, we focused on the period of TRFs from −100 ms to 400 ms but showed the TRFs from −200 ms to 500 ms. The EEG signals and the amplitude envelopes of music pieces were not filtered or decomposed into different frequency bands. We fixed the lambda at 0.1 for calculating TRFs across all the conditions and tempi and hence the differences of TRFs across conditions should not be because of different lambda values. After calculating TRF for each music piece, we averaged TFRs over 10 pieces of one condition for each participant and conducted statistic tests on the mean TFRs across different conditions. The TRFs were calculated using the Multivariate Temporal Response Function (mTRF) Toolbox (Crosse et al., 2016).

The fixed lambda value of 0.1 was decided empirically. The lambda value decides how signals are temporally smoothed during TRF calculation and makes sure that the encoding model does not overfit to noise: the larger the lambda value is, the signals are smoothed to a larger extent temporally – the lambda functions similarly as the cutoff frequency of a low-pass filter. If zero is chosen for the lambda, the inversion of matrices cannot be implemented as sometimes the covariance of neural responses recorded here is not fully ranked. Therefore, we chose 0.1 as the smallest value of the lambda to make sure that the inversion can be implemented, and the signals were not smoothed seriously. We believe that the choice of the lambda value should not affect our results, as we compared TRFs across the conditions instead of drawing conclusions from the absolute TRF values and the choice of the lambda value exerted an equal influence on TRFs of all the conditions.

#### Temporal modulation of EEG power

Here we aim to investigate musical phrase segmentation beyond the level of the beat structure. The lengths of the music phrases ranged from ∼7 seconds (8 beats per music phrase at 65 bpm) to ∼5 seconds (8 beats per music phrase at 85 bpm), which corresponded to an ultra-low EEG frequency range of ∼0.13 Hz to ∼0.18 Hz. This ultra-low EEG frequency range has proved to be challenging to analyze because of the 1/f shape of power spectrum of EEG recording and hence a low signal-to-noise ratio at the ultra-low frequency range. Another fact is that the time-frequency analysis requires an ultra-long temporal window for such frequency ranges (i.e. two cycles for 0.13 Hz, which is ∼15 seconds). Previous studies have not yet shown a meaningful and robust neural signature within this range. It is indeed challenging to directly observe temporal dynamics of musical phrase segmentation and its corresponding frequency components in EEG signals.

To circumvent the challenge of analyzing the ultra-low frequencies of EEG signals and to directly observe temporal dynamics of musical phrase segmentation, we resorted to a different strategy: as we observed the boundary effects of neural responses to the boundary beats of musical phrases (Fig. 2G), the musical phrasal structures likely modulated the power of neural responses to each beat. The beat rates of the music pieces were above 1 Hz, at which neural signals can be well recorded by EEG, and the analyses above showed robust beat tracking (Fig. 2). Therefore, we chose to measure the temporal modulation of EEG power at the beat rate to investigate musical phrasal tracking. If listeners do track musical phrases and the neural responses to beats are modulated by musical structures, we should be able to observe neural signatures reflecting musical phrase segmentation through temporal modulation of EEG power at the beat rates. This procedure is akin to recovering low-frequency amplitude envelopes from high-frequency carriers in speech signal processing (Teng et al., 2019).

We first conducted a time-frequency analysis on EEG signals to derive power spectrograms of neural responses to the music pieces. The single-trial data derived from MCCA were transformed using the Morlet wavelets embedded in the Fieldtrip toolbox, with a frequency range from 1 to 35 Hz in steps of 1 Hz and a temporal range from 3 seconds before the onset of music pieces to 3 seconds after the offset of music pieces in steps of 100 ms. To balance spectral and temporal resolution of the time-frequency transformation, from 1 to 35 Hz, the window length increased linearly from 3 cycles to 10 cycles. Power (squared absolute value) was extracted from the wavelet transform output at each time-frequency point. Moreover, the power values for each trial were normalized by dividing the mean power value over the baseline range (−1.5 s ∼ −.5 s) and taking logarithms with base 10, and then was converted into values with unit of decibel by multiplying by 10.

We next quantified temporal modulation spectra of EEG power to examine at which frequency the EEG power was modulated and whether the salient modulation frequencies were related to the frequency range of musical phrasal structures. This procedure is similar to the calculations of acoustic modulation spectra of the music pieces (Fig. 1C). We calculated the modulation spectrum of EEG power at each frequency using FFT with zeropadding of 20000 points and took the absolute value of each frequency point, so that the spectral resolution of the modulation spectra was 0.005 Hz. The high spectral resolution guaranteed to resolve different music phrase rates (8 beats per music phrase: 0.1375 Hz at 65 bpm; 0.1562 Hz at 75 bpm; 0.1771 Hz at 85 bpm). We then normalized the modulation spectra of each trial by dividing the modulation spectrum of each trial by the length of its corresponding music piece. As the music pieces of each condition had different lengths, they could not be summed together in the temporal domain, we chose to sum the modulation spectra (complex numbers) of 10 pieces within each condition in the spectral domain. This spectral summation on complex number reserved phase information, and the shared frequency components with similar onset phases across the 10 pieces in a condition were emphasized. After the spectral summation, we took the absolute value of each frequency point to derive the modulation spectrum for each condition. The above procedure resulted in a two-dimensional modulation spectrum for each condition, with one dimension as the frequency of EEG power and the other dimension as the frequency of temporal modulation of EEG power. It should not matter for the following statistic tests in this case whether we summed the modulation spectra or averaged the modulation spectra over the music pieces, as both summation and averaging are linear operations.

To examine how the musical phrase segmentation varied with the tempo of music pieces, we averaged over three reversing conditions within each tempo and showed the modulation spectra of three tempi, respectively (Fig. 3A). If the neural signatures of musical phrase segmentation varied with the tempo, this helped validate the observed neural signatures. We would also like to check whether other frequency ranges of EEG power (i.e., the alpha band, 8 – 12 Hz, and the beta band, 13 - 30 Hz) showed signatures of musical phrase segmentation, as the previous literature reported an important role of the neural beta band in music processing (Doelling and Poeppel, 2015; Fujioka et al., 2015).

We next focused on the frequencies of EEG power at the beat rates of three tempi, as we only observed salient modulations of EEG power around 1 Hz (Fig. 3A), which was close to the beat rates of three selected tempi. We conducted once again the time-frequency transformations on the EEG signals, but only at the beat rates, 1.1001 Hz (65 bpm), 1.25 Hz (75 bpm), and 1.4165 Hz (85 bpm). The window length in the time-frequency transformations was 3 cycles. Other procedures and parameters were the same as the above analyses of calculating the EEG power and the modulation spectrum of each tempo and each reversing condition. We selected the third music piece of the original condition (BWV267), which had the longest duration, and showed the temporal dynamics of EEG power at the beat rate of each tempo (Fig. 3B). This directly showed how the EEG power was modulated by the musical phrasal structures. For each participant, we summed the modulation spectra of 10 pieces within a condition in the spectral domain and conducted statistics on the summed modulation spectrum of each condition (Fig. 3C).

#### Surrogate test on modulation spectra of EEG power

The modulation magnitudes of neural signals of musical phrase segmentation cannot be directly compared between different tempi, as the music pieces of different tempi had different lengths and the tempi biased estimation of the modulation spectra reflecting the musical phrase segmentation– frequency components in the lower frequency range tend to have larger strength because of the 1/f nature of neurophysiological signals. Therefore, individual baselines need to be constructed for the modulation spectra of different tempi; then the modulation spectra can be normalized according to the baselines within each tempo. Therefore, we conducted surrogate tests in the spectral domain to derive a null distribution of the modulation spectrum for each condition and each tempo. This served two purposes – we can directly test the significant frequency ranges of musical phrase segmentation using the null distributions and then normalize the modulation spectra to do comparisons across conditions.

The null hypothesis of the surrogate test here is that the neural signals did not track or phase-lock to the musical phrasal structure, and hence the sum of modulation spectra across 10 pieces of each condition should not differ from the surrogate modulation spectra derived from jittering the temporal alignment between each of the 10 music pieces and its neural recording temporally. However, it is not appropriate to conduct this surrogate test in the time domain. As the lengths of musical phases (> 5 seconds) were much longer than the pre- and post-stimulus periods (3 seconds), temporal surrogation cannot efficiently disrupt the temporal correspondence between neural signals and musical phrasal structures - the jittering can only happen within a cycle of musical phrase. More importantly, the tempi here still affect the procedure of surrogation – for example, jittering within a temporal range of 6 seconds at 85 bpm has a different jittering effect from the same jittering range at 65 bpm because of different lengths of musical phrases at the two tempi. Therefore, we conducted the surrogate test in the spectral domain after Fourier transformation of EEG power.

After calculating the modulation spectrum of the EEG power of each music piece, we multiplied the modulation spectrum (complex numbers) with a complex number that has a norm of 1 but a randomly generated phase from 0 to 2 * pi. This multiplication reset the onset phase of the EEG power of each music piece so that the EEG power of each music piece now started with a random phase, whereas the amplitude magnitude of the modulation spectrum was kept intact. We then derived the summed modulation spectrum of the surrogated data for each tempo and each condition. By doing this, the 10 pieces within a condition all had different onset phases and therefore the temporal components locked to the musical phrasal structures should be averaged out or severely smoothed. This surrogating procedure on phase in the spectral domain was invariant to different tempi, as the phase values were normalized to different tempi. We repeated this procedure 1000 times and derived a null distribution of modulation spectra for each condition and each tempo and for each participant.

We averaged the modulation spectra of each surrogation across participants and derived a null distribution of the group-averaged modulation spectrum for each condition and each tempo. We chose a one-side alpha level of 0.01 as the significant threshold for each condition and each tempo (dashed lines in Fig. 3C). The frequency ranges where the empirical modulation spectra are above this threshold can be considered as the frequency ranges showing robust effects of musical phrase segmentation. To derive one frequency range across three conditions for one tempo so that the frequency range selected is unbiased to each condition for the following analysis, we chose the lower bound and the upper bound of the significant frequencies among all the three conditions and used them to define one frequency range for all the three conditions. The significant frequency ranges are as following: at 65 bpm, 0.125 – 0.15 Hz; at 75 bpm, 0.14 – 0.175 Hz; at 85 bpm, 0.165 – 0.2 Hz.

#### Normalization of modulation spectra of EEG power

From the surrogate tests above, we obtained the null distributions for the modulation spectra of EEG power. As mentioned, using the null distributions we could normalize the raw modulation spectra that were biased because of different tempi of music pieces. Here, we calculated the mean of the null distribution for each condition and each tempo and subtracted out the mean of the null distribution from the raw modulation spectrum.

#### Temporal response function of EEG power envelope with phrasal boundaries as regressor

In the above spectral analysis, we characterized the frequency components reflecting musical phrase segmentation. What are the temporal trajectories of musical phrase segmentation? Is the musical phrase segmentation a result of neural signals locking to the onset beat or to the offset beat of a musical phrase? The spectral analyses cannot reveal such temporal information. Therefore, we calculated TRFs of EEG power using the phrasal boundaries of music pieces as the regressors.

The TRFs of EEG power envelopes were calculated in a similar way as in the calculation of TRF of boundary beats of musical phrases. We marked the boundaries between two phrases in each music piece. For example, there are 7 phrases in the music piece of BWV153/1 (see Table S1, Piece Number 1) and hence there are 6 phrase boundaries; these 6 phrase boundaries served as a regressor for the EEG power envelope to derive a TRF for this piece. After deriving the EEG power envelopes at each tempo, we calculated TRF for each music piece from 1 second after stimulus onset to 1 second before stimulus offset. The length of TRF estimation was 6 seconds, with 3 seconds before the phrase boundary and 3 seconds after. We then averaged TRFs of 10 pieces within a condition of each tempo. The lambda was set as 0, which indicated that no smoothing was applied and the calculation of TRF was equal to a reverse correlation.

The TRFs of EEG power envelope are shown in Fig. 3E. It can be seen that the peak latencies for different conditions differ. To determine the latency of each condition and to find the peak time point, we fitted each group-averaged TRF using a one-term Gaussian model, whose center point determined the peak latency of each group-averaged TRF. The model fits were plotted as inserts in Fig. S2C.

#### Mutual information between music criteria

We defined musical phrasal boundaries according to the notifications of music scores, but this a priori definition of the musical phrasal structure may not be best to explain the musical phrase segmentation we observed here. As the different reversing manipulations broke musical structures to different extents, by calculating MI between different criteria, we could see how coherent the four musical criteria are with each other. This MI measurement helped answer whether the decreased musical phrase segmentation of one condition could be because of the incongruence between musical criteria caused by the corresponding reversing manipulation – congruent cues of different musical criteria collectively help listeners to parse the music pieces into high-level musical units beyond note and beat, whereas the incongruence among the musical criteria causes confusion to music stream segmentation. The congruence between music criteria may help explain why the participants better tracked musical phrases in one condition than the others.

We first calculated MI between different musical criteria. We set the boundary beat marked by a musical criterion to 1 and the other beats to 0. Hence, we had a vector of ‘0’ and ‘1’ for each music piece according to one music criterion. The time unit of this vector is the number of beats. We show an example of one criterion vector in Fig. S5A. We then computed the normalized MI between the four musical criterion vectors for each music piece. The normalized MI was calculated as the following with x and y representing two different criterion vectors:

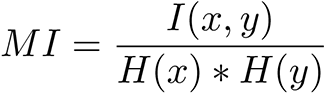

After deriving the MI between different criteria for each music piece, we averaged the music pieces within each condition and used the averaged MI as a measure for the MI results between two musical criteria in one condition. The results are plotted in Fig. S5B. On the other hand, to obtain a measure for the congruence between the four criteria for each music piece, we averaged over all the MI results between the four criteria for each music piece and used the averaged MI of each piece as the congruence measure. If all the criteria are highly coherent with each other, the averaged MI is close to 1. Otherwise, the average MI is close to 0. We plotted the congruence between the musical criteria for each piece in Fig. S4C.

#### Temporal response function of EEG power envelope with musical criteria as regressor

Following the same procedures used in *Temporal response function of EEG power envelope with phrasal boundaries as regressor*, we calculated TRFs of EEG power using the segment boundaries defined by different musical criteria as the regressors, and showed the results in Fig. S4D.

#### Phrasal phase precession and phrase segmentation at group-average level

The original music pieces are well structured, and a listener can readily extract the temporal regularity of the phrase structures and predict the unfolding of the musical pieces. In contrast, the reversed conditions may preserve the salient onset and offset beats of musical phrases or expected chord progressions but reduce the structural cues to the phrases, so listeners experience difficulties to predict the unfolding of the pieces. To test this conjecture, we tried to quantify phase precession of neural responses on the level of musical phrases – *phrasal phase precession (PPP)*, as phase precession of neural signals has been argued to reflect the prediction of future events (Jensen and Lisman, 1996; Lisman, 2005). The rationale here is that, if listeners can predict the incoming musical contents, the neural phase should proceed faster as the music unfolds. On the other hand, if listeners cannot predict the incoming music contents but passively respond to or are solely driven by the acoustic events in music, the neural phases would proceed at a constant speed, as the speed/tempo of the acoustic events is constant in each music piece.

To quantify phrasal phase precession, we need to conduct our analysis on the neural response to each music piece at a single-trial level. However, we conducted EEG recording here and the EEG recording at a single-trial level often has a low signal-to-noise ratio; usually at least tens to hundreds of repetitions are needed for analyzing neural responses to a single stimulus. To resolve this issue, we took another strategy for quantifying phase precession for each music piece – we first averaged each trial across 29 participants and then conducted our analyses on the group-averaged trial. Although such strategy prevented us from probing individual differences, the process of the group averaging did aid in emphasizing the neural components induced by the music pieces while averaging out neural signals unrelated to the music listening. If a music piece is predictable on the level of musical phrase segmentation, the phrasal phase precession should be observed for all the participants to various degrees. Therefore, the phase precession quantified from the group-averaged trials could collectively indicate that the participants predict the unfolding of each music piece as well as reflect the predictability of each music piece.

The above analyses had extracted the EEG power envelopes reflecting the musical phrase segmentation and determined the significant frequency ranges of musical phrase segmentation at each tempo. We averaged the power envelopes of each music piece across 29 participants and employed a two-pass second-order Butterworth filter embedded in the Fieldtrip toolbox to filter the group-averaged envelopes, using the significant frequency range of each tempo as the cutoff frequencies. The phases of the averaged power envelope of each music piece were extracted by applying the Hilbert transformation on the filtered signal and taking the angle at each time point. We showed in Fig. 4A (left panel) the phase series of three conditions of the music piece, BWV267, at 75 bpm, as this piece has the largest length. We also plotted the phase values at the phrase boundaries in Fig. 4A (right panel) so that it can be clearly seen that the proceeding speed of the neural phases was increasing in the original and global reversal conditions as the music piece unfolded.

We constructed an index, *phrasal phase precession index (PPPi)*, to quantify phrasal phase precession over each music piece. If there is no phase precession, the neural phase series would proceed in a step of 2 * pi over every musical phrase. This is to say that, if we fit a line between the musical phrasal boundaries and their corresponding neural phase values, the slope will be exactly 2 * pi if there is no phase precession. In contrast, if the neural phase series is accelerating as the music piece is unfolding, this line will be steeper and the slope will be larger than 2 * pi; if the phase series is slowing down, the slope will be smaller than 2 * pi. Therefore, the differences between the slope of the fitted line and 2 * pi can represent whether there is phase precession and to what extent the phase series is accelerating or slowing down. The differences here were termed as PPPi. We provide an example in Fig. 4C, where we show the lines fitted between the music phrase boundaries and their corresponding neural phase values in three reversing conditions at 75 bpm.

The peak frequencies of modulation spectra of EEG power correlate with PPPi, as the phase precession can be considered as frequency modulation or phase modulation – the frequency of musical phrase segmentation is modulated along the time – and the mean frequency over the whole music piece becomes larger than the phrase rate if the phrasal phase precession happens. Therefore, here, we also quantified the peak frequencies of the modulation spectrum of EEG power within the significant phrase segmentation range for each music piece, so that the results of PPPi can be further validated by a different index.

To detect the peak frequency within the significant phrase segmentation range derived from the above (*Surrogate test on modulation spectra of EEG power*), we first looked for the frequency point of the largest modulation amplitude of each group-averaged trial for each music piece within the significant phrase segmentation frequency range, and then we calculated the differences of modulation amplitude between its adjacent frequency points. If the differences indicated that this frequency point sat at a concave function (negative difference on the left and positive difference on the right), we considered this frequency point to be the peak frequency of the modulation spectrum for the music piece. Otherwise, it meant that we did not find a peak frequency – either the curve of modulation spectrum within the significant phrase segmentation frequency range is not concave or the peak frequency is at the boundary of the significant range. The results are shown in Fig. S6.

#### Phrasal phase precession at the level of individual participants

As the significant frequency ranges of the phrase segmentation defined in Fig. 3C was derived from the group-averaged data, the frequency ranges were not applicable to the individual participant’s data. Therefore, for each participant, we calculated PPPi of each music piece at each tempo (from left to right) for each condition using the bandwidth from ± 0.05 octave to ± 0.14 octave (from top panel to bottom panel), so that the effect of the bandwidth on the PPPi calculation can be examined. For example, we chose a bandwidth of ± 0.05 octave at the phrase rate of 0.138 Hz and hence the frequency range for calculating PPPi in this case was from 0.1333 Hz to 0.1429 Hz. The procedures of calculating PPPi were the same as in *Phrasal phase precession and phrase segmentation at group-average level*, but were conducted on the individual data instead of the group-averaged data.

#### EEG topographies and source localization

We re-conducted the analyses of Cacoh, TRF, and phrase segmentation on each EEG electrode and in the source space to show topographies and source localization results (Fig. S2&S4), which complemented the above analyses based on MCCA components. Note that MCCA extracted the neural components key to music listening and overcame variations of individual participants’ head sizes and EEG cap positions during EEG recording. In contrast, the analyses on the EEG electrodes and in the source space failed to do so. Therefore, we did not conduct statistical tests in the electrode space and the source space but only visualized the neural signatures.

We conducted the same analysis procedures of Cacoh, TRF, and phrase segmentation on the preprocessed data of each electrode before running MCCA. As the analyses on the MCCA components already tested the effects of different conditions, we selectively showed Cacoh values at the beat rate and the note rate of the three tempi (Fig. S3A), and the modulation strength of phrase segmentation of the three reversal conditions (Fig. S3B).

We next conducted EEG source reconstruction and localized Cacoh and modulation strength of phrase segmentation. As no structural T1-weighted MRI scan was acquired, we used a template boundary element model (BEM) of a resolution of 5 mm provided by EEGLAB (Delorme and Makeig, 2004) as the head model. The BEM model was composed of three 3-D surfaces (skin, skull, cortex) extracted from the MNI (Montreal Neurological Institute) canonical template brain. The source reconstructions were done by Minimum Norm Estimation (MNE) implemented in the Fieldtrip toolbox 20181024, and the covariance matrix was calculated using the EEG data of two seconds before the onsets of all the 90 music pieces. The forward solution was estimated from a source space of 8196 activity points; the inverse solution was calculated from the forward solution. We did not scan a head shape nor measure EEG cap position for each participant, so the precision of the source location results was severely limited. Nevertheless, the source localization can be used to provide guidance for explaining our findings. After projecting the preprocessed EEG data to the source space, we conducted the analyses procedures of Cacoh and phrase segmentation on each virtual channel (each vortex). We did not conduct *Surrogate test on modulation spectra of EEG power* in the source space, so the modulation strength of phrase segmentation was of raw values.

### DATA AND CODE AVAILABILITY

The music materials, group-averaged EEG data, and all the analysis codes have been deposited in the OSF folder (https://osf.io/vtgse/). The preprocessed individual EEG datasets will be uploaded to the OSF folder and be fully available by 31^st^, January, 2022, as the authors and their collaborators are developing further studies based on the EEG datasets of this study. Readers can contact the Lead Contact, Xiangbin Teng, to request the raw and the preprocessed EEG datasets for validation and replication of the current study.

## Acknowledgement

We thank Johannes Messerschmidt, Dominik Thiele, Claudia Lehr and Cornelius Abel for their technical support; Johannes Messerschmidt for his assistance with data collection; Cecilia Musci and Lea Fink for their assistance in preparing music materials. We thank Yi Du, Molly Henry, and Andrew Chang for their comments on a previous version of the manuscript. This work was supported by the Max-Planck-Society.

## Supplementary Information

**Fig. S1.**
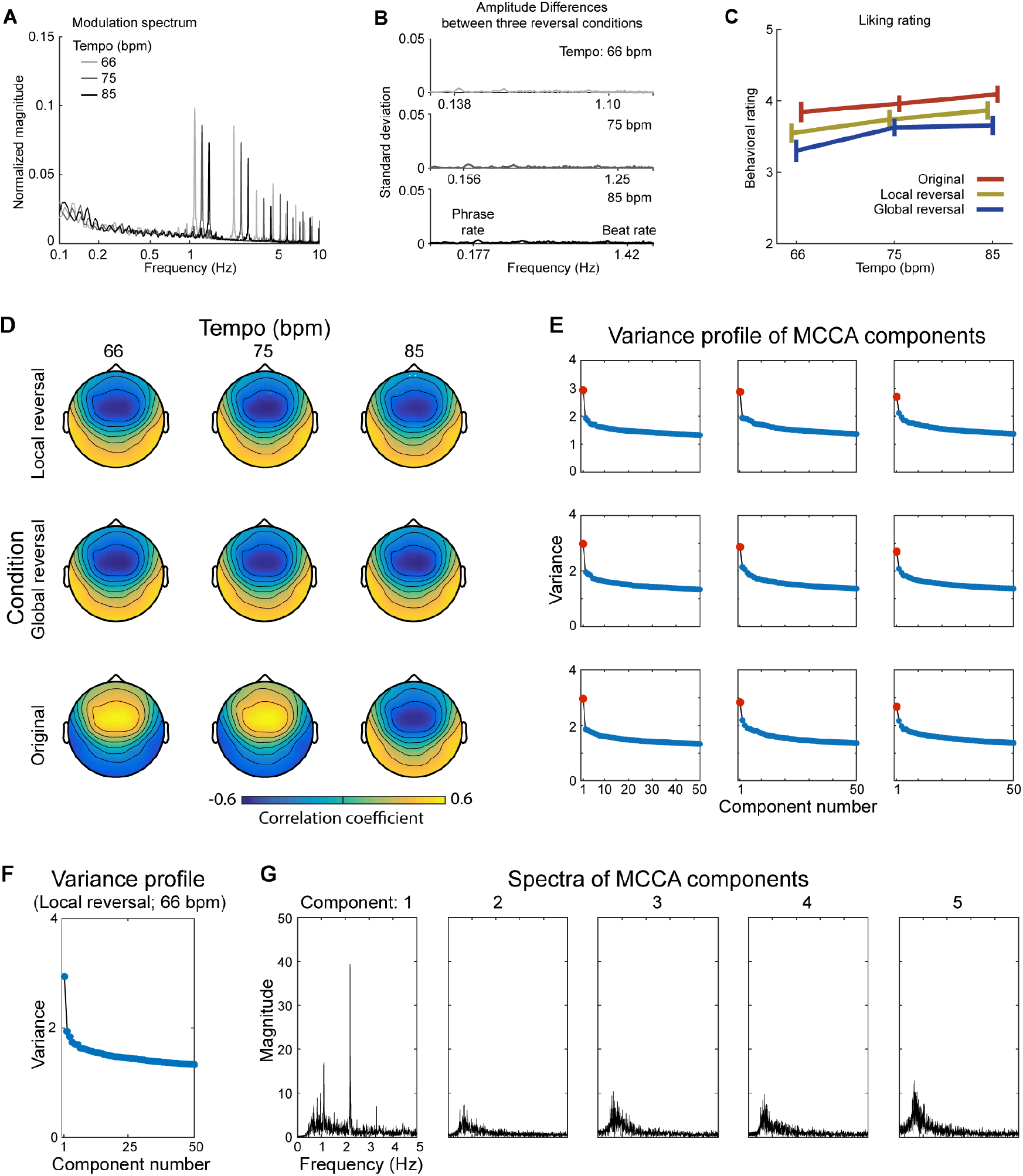
Acoustic analyses, behavioral rating, and MCCA procedure, related to Fig. 1&2. (A) Acoustic analyses. We extracted the amplitude envelopes of the music pieces and calculated the modulation spectra averaged over 10 music pieces of each tempo in each reversal condition. The modulation spectra of the three reversal conditions highly overlapped and hence were plotted together. The fundamental frequency of each tempo represents the beat rate. (B) The standard deviation across three reversal conditions. In terms of amplitude fluctuations, the three conditions are comparable at each tempo. This demonstrates that the acoustic properties of the music pieces do not differ and should not contribute to the difference of phrasal segmentation between the reversal conditions. (C) Liking rating. The participants rated how much they liked each piece from 1 to 6. We conducted a two-way repeated-measure ANOVA (rmANOVA) with Condition and Tempo as the main factors, and found that both Tempo (F(2,62) = 7.80, p = 0.001, ηp2 = 0.201) and Condition (F(2,62) = 26.69, p < 0.001, ηp2 = 0.463) affected the participants’ liking ratings, but the interaction was not significant (F(4,124) = 1.20, p = 0.314, ηp2 = 0.037). The listeners assigned higher liking ratings to faster pieces; the Original condition was rated higher than the other two conditions (Original > Global reversal: t(31) = 6.76, p < 0.001; Original > Local reversal: t(31) = 3.54, p = 0.001); the Global reversal received the lowest liking rating (Local reversal > Global reversal: t(31) = 4.55, p < 0.001). Adjusted False Discovery Rate (FDR) was used for multiple comparison correction. (D) Topographies of correlation between the first MCCA component and each EEG channel. We calculated the Pearson correlation of neural signals between the first MCCA component and each EEG channel for each music piece. We then averaged the 10 music pieces of each tempo in one condition and plotted the topography. As the principle component analysis sometimes reverses the sign of a component, all the conditions did not have the same positive or negative signs. (E) MCCA components in each condition. We show 50 MCCA components in each condition. The first component explained a disproportionally large variance. This supports our choice of only extracting and proceeding with the first component for analysis. (F) MCCA components of the Local reversal condition at the tempo of 65 bpm. (G) Amplitude spectra of first 5 MCCA components of the Local reversal condition at the tempo of 65 bpm. We calculated the amplitude spectrum of each MCCA component in (F) to examine the spectral component of each component. The spectrum of the first component shows amplitude peaks corresponding to the beat rate and the note rate (the first harmonic of the beat rate). This further strengthens our choice of analysis – specifically, the first MCCA component, but not other components, contained the neural signals induced by beat and note structures in the music pieces. Furthermore, this echoes the topographies in (D), showing that the topographies reflected auditory responses to beats and notes.

**Fig. S2.**
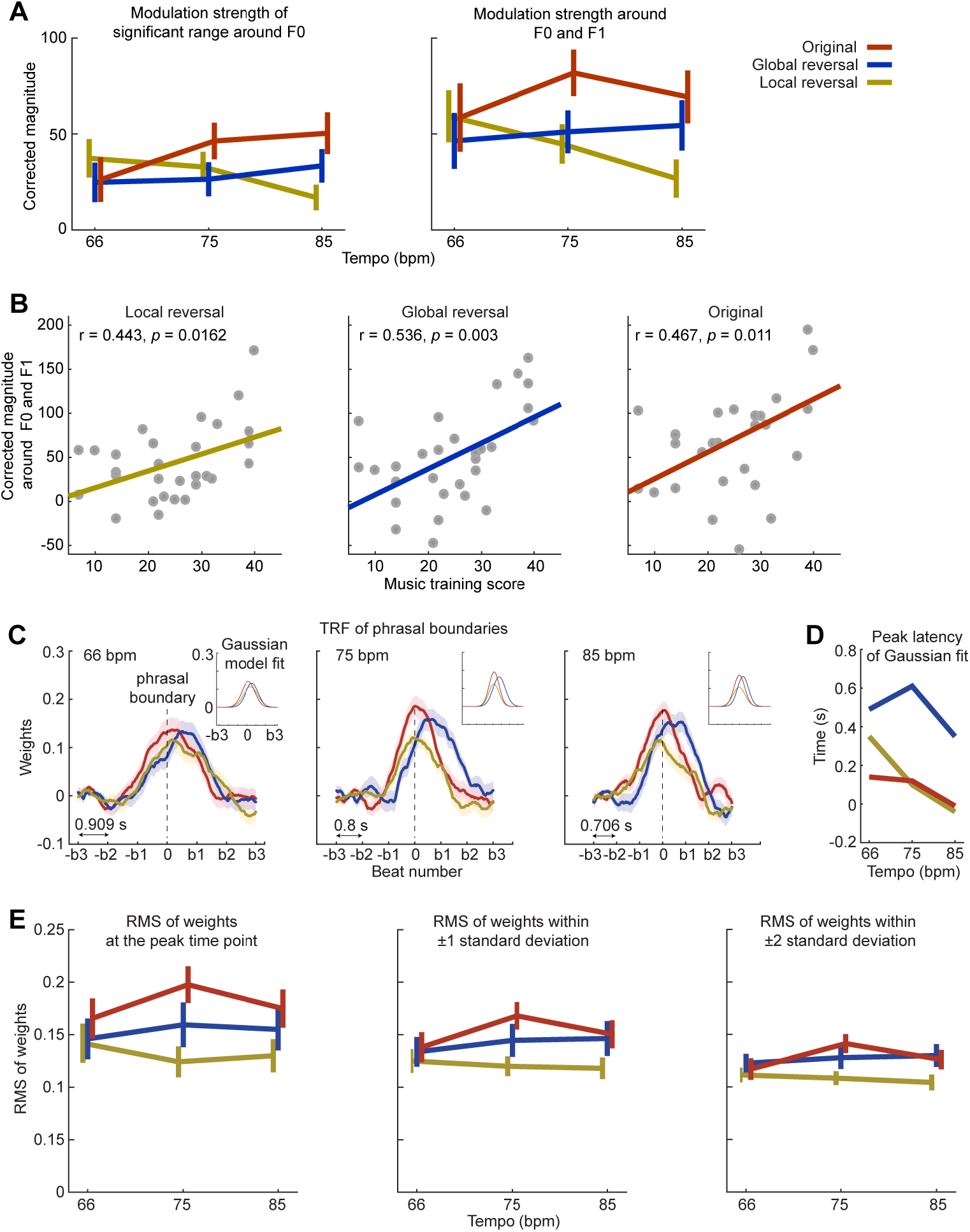
Modulation magnitude around F0 and F1, and RMS of TRFs, related to Fig. 3. (A) Modulation magnitude around F0 and F1. We averaged the corrected modulation magnitude within the significant frequency ranges identified in Fig. 4C around the fundamental frequency of the phrase tracking, F0 (left panel) and conducted a two-way rmANOVA (Tempo × Condition). We did not find any significant main effect nor interactions (p > 0.05). This is likely due to the fact that the neural signals locking to phrasal structures are of complex waveforms, instead of sinusoid waves, and the narrow frequency range around the phrase rate cannot capture all the relevant neural components. We then averaged the magnitude over the significant frequency ranges around F0 of the phrase tracking and its first harmonic (F1) (right panel), and the two-way rmANOVA showed a significant main effect of Condition (F(2,56) = 3.66, p = 0.032, ηp2 = 0.115), but not of Tempo (F(2,56) = 0.66, p = 0.521, ηp2 = 0.023) or the interaction (F(4,112) = 1.48, p = 0.214, ηp2 = 0.050). In the post-hoc test, after FDR correction, we did not find any differences between the reversal conditions. However, before FDR correction, the Original reversal is significantly larger than the Local reversal (t(28) = 2.36, p = 0.026). The error bars represent ±1 standard error of mean over the participants. (B) Correlation between music training score and EEG power modulation of each reversal condition within the significant frequency ranges around both F0 and F1. The results are consistent with Fig. 3D. (C) TRFs of phrasal boundaries and the Gaussian model fits. The TRFs are the same as in Fig. 3E and we fitted Gaussian models to the group averaged TRFs (insert in each panel) and derived the latencies of TRF peaks and the standard deviations of TRFs (see Methods). The shaded areas represent ±1 standard error of mean over the participants. (D) Peak latency of TRF. Interestingly, the Global reversal had a peak latency much larger than the other two reversal conditions, which suggests that the reversing procedure changed the phrasal structures so that the listeners segmented the high-level musical structures differently. This observation was further examined in Fig. S6. (E) We calculated RMS of TRFs at the peak time point (left panel), within ±1 standard deviation (middle panel), and within ±2 standard deviation (right panel), which were determined by the Gaussian fits in (C). We tested the significance of the differences of RMS between the conditions by conducting a two-way rmANOVA (Tempo × Condition) on the RMS calculated using each time range. When the peak points were used, we found a significant main effect of Condition (F(2,56) = 5.73, p = 0.005, ηp2 = 0.170), but not of Tempo (F(2,56) = .31, p = 0.732, ηp2 = 0.011) nor the interaction (F(4,112) = 0.68, p = 0.608, ηp2 = 0.024). When ± 1 standard deviations were used, we found a significant main effect of Condition (F(2,56) = 4.49, p = .016, ηp2 = 0.138), but not of Tempo (F(2,56) = 0.93, p = .399, ηp2 = 0.032) or of interaction (F(4,112) = 0.68, p = 0.607, ηp2 = 0.024). The post-hoc test shows that the Original condition is significantly larger than the Local reversal (t(28) = 2.96, p = .018, FDR corrected). When ± 2 standard deviations were used, we found a significant main effect of Condition (F(2,56) = 4.72, p = 0.013, ηp2 = 0.144), but not of Tempo (F(2,56) = .92, p = 0.404, ηp2 = 0.032) nor the interaction (F(4,112) = 0.95, p = 0.439, ηp2 = 0.033). The TRF results are consistent with the spectral analysis using both F0 and F1 depicted in (A, right panel). The error bars represent ±1 standard error of mean over the participants.

**Fig. S3.**
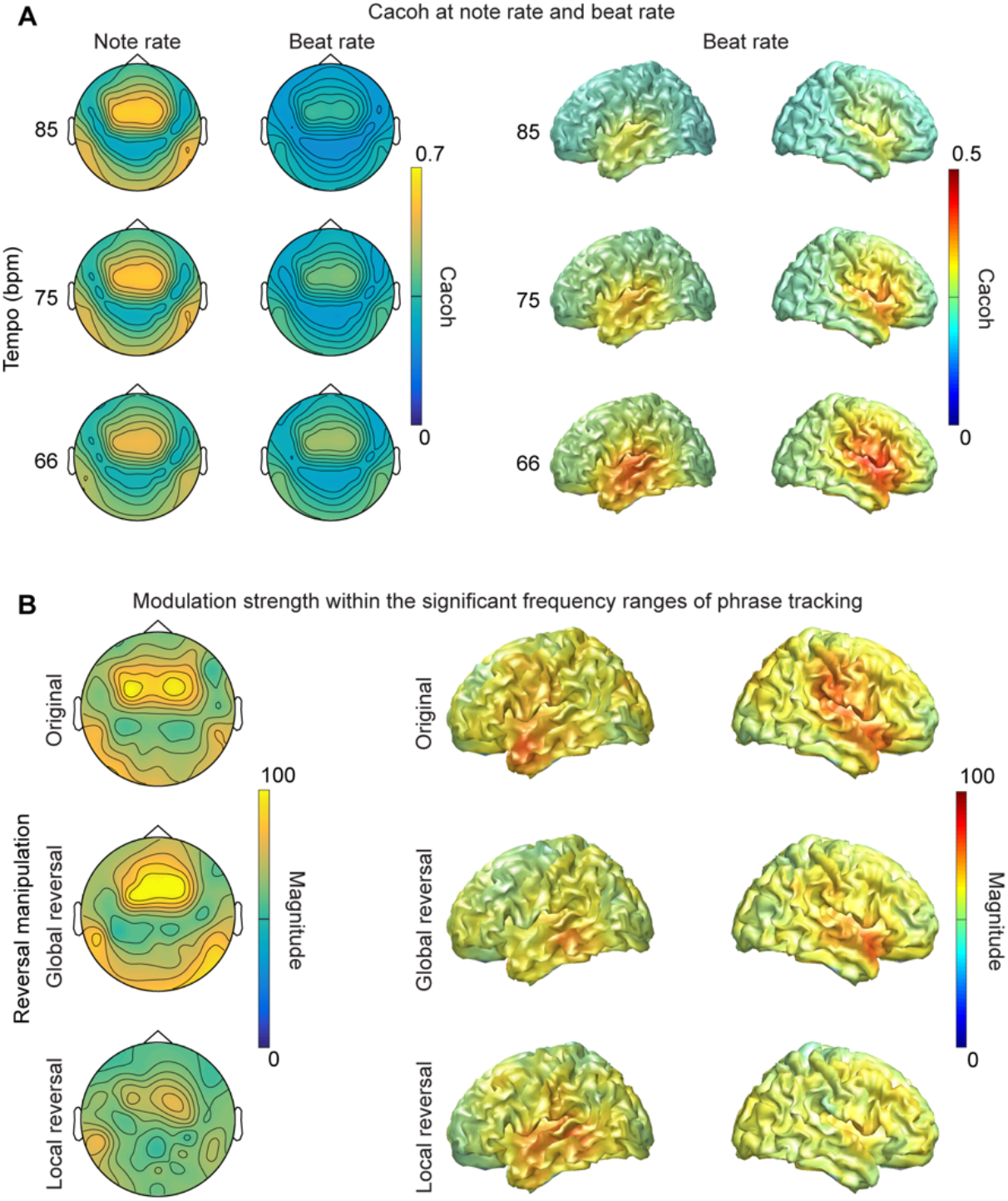
Topographies and source localization plots of Cacoh and phrasal segmentation, related to Fig.2&3. (A) Cacoh at note and beat rates. We averaged Cacoh values at the note and the beat rates over three reversal conditions, as the main effect of Condition was not significant (p < 0.05) (see Fig. 2), and show topographies of different tempi in the left panel and source localizations in the right panel. The same analysis procedures were conducted at the level of electrodes and in the source space as in Fig. 2. (B) Phrasal segmentation. We averaged modulation magnitudes within the significant frequency range at F0 of the phrasal rate identified in Fig. 3C, and show topographies of different reversal conditions in the left panel and source localizations in the right panel. The modulation magnitudes were of raw modulation strength and were not normalized. Other than that, the same analysis procedures were conducted at the level of electrodes and in the source space as in Fig. 3C.

**Fig. S4.**
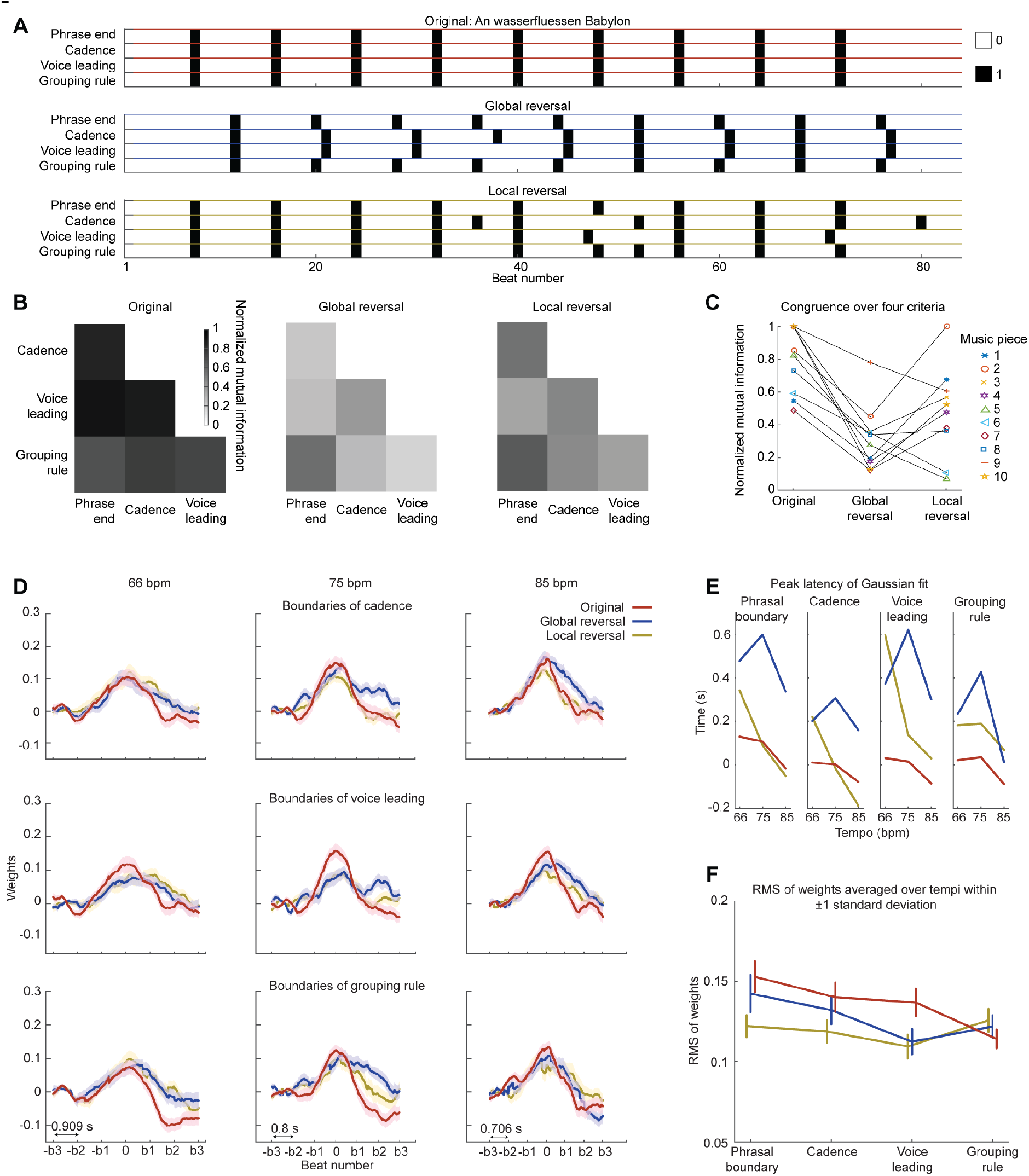
Mutual information (MI) analyses on musical criteria and TRFs, related to Fig. 3. (A) Musical criteria for an example music piece of the three reversal conditions. Four criteria were used here – pre-defined phrase end, cadence, voice leading, and grouping rule. Each beat was treated as a temporal unit and the structural boundaries determined by the criteria were marked on the beat number. The black-filed boxes indicate the beats representing the structural boundaries. The matrices for the following calculation of mutual information were created by marking the black boxes as ‘1’ and the other beats (empty areas/boxes) as ‘0’. (B) MI between different musical criteria. We quantified similarity between different musical criteria by calculating normalized mutual information between each pair of musical criteria (see Methods). In the Original condition, the musical criteria are highly coherent; it is not the case in the Global reversal and Local reversal conditions. (C) Congruence over four musical criteria for each music piece. We averaged the MI values over all the pairs of musical criteria for each music piece and used the averaged MI value as an index for the congruence over the four musical criteria of a music piece. It can be seen that most of the music pieces in the Global reversal condition have low congruence. The congruence pattern here is not consistent with the music phrasal segmentation (Fig. 3&S2), as the segmentation strength in the Global reversal condition is between the Original condition and the Local reversal condition. (D) We calculated TRFs of EEG power using the boundaries defined by four musical criteria as the regressor. This analysis procedure is the same as in Fig. 3E. The x axis is marked in the number of beats; the double arrow line indicates the length of the beat at each tempo. We fitted Gaussian models to the group-averaged TRFs and derived the latencies of TRF peaks and the standard deviations of TRFs. The shaded areas represent ±1 standard error of mean over the participants. (E) Peak latency of TRFs. The group-averaged TRF of the Global reversal calculated using the phrasal boundaries had a large peak latency, but the latency decreased when the TRF was calculated using the cadence and grouping rule, which suggests that the phrasal segmentation in the Global reversal condition probably locked to the cadence or the grouping rule. See Fig. S6 for more information. (F) RMS of TRFs within ±1 standard deviation. We first calculated RMS of TRFs from different musical criteria within ± one standard deviation for each tempo and each condition, and conducted a three-way rmANOVA (Criteria * Condition * Tempo). We did not find a significant main effect of Tempo (F(2,56) = 1.91, p = 0.158, ηp2 = 0.064); the interaction is not significant between Tempo and Criteria (F(6,168) = 1.98, p = 0.071, ηp2 = 0.066) nor between Tempo and Condition (F(4,112) = 1.42, p = 0.231, ηp2 = 0.048). The interaction between the three factors is not significant (F(12,336) = 0.404, p = 0.962, ηp2 = 0.014). We then averaged RMS across the tempi and conducted a two-way rmANOVA (Criteria * Condition). We found a significant main effect of Criteria (F(3,87) = 0.404, p < 0.001, ηp2 = 0.196) and a significant interaction (F(6,174) = 5.50, p < 0.001, ηp2 = 0.159); the main effect of Condition is not significant (F(2,58) = 3.05, p = 0.055, ηp2 = 0.095). We tested the difference between each pair of musical criteria to further examine the main effect of Criteria. After FDR correction, the RMS of the phrase boundary is significantly larger than the voice leading (t(28) = 4.37, p < 0.001) and grouping rule (t(28) = 2.78, p = 0.019), but not than the cadence (t(28) = 2.16, p = 0.059). The RMSs of the cadence, the voice leading and grouping rule are not significantly different from each other (p > 0.05, FDR correction). The error bars represent ±1 standard error of mean over the participants. To explore the interaction between Condition and Criteria, we conducted a one-way rmANOVA (Condition) on each musical criterion. After FDR correction, we found a significant main effect of Condition for the phrasal boundary (F(2,58) = 4.80, p = 0.023, ηp2 = 0.142) and the voice leading (F(2,58) = 6.50, p = 0.012, ηp2 = 0.183), but not for the cadence (F(2,58) = 3.38, p = 0.055, ηp2 = 0.104) nor for the grouping rule (F(2,58) = 0.819, p = 0.446, ηp2 = 0.027). In summary, the phrasal boundary better explained the temporal modulation of EEG power than the other three musical criteria. But the cadence also showed comparable capacity of explaining the phrasal segmentation results. This suggests that the phrasal segmentation observed was partly driven by the cadence of the music pieces, but cannot be fully attributed to the cadence structures in the music pieces.

**Fig. S5.**
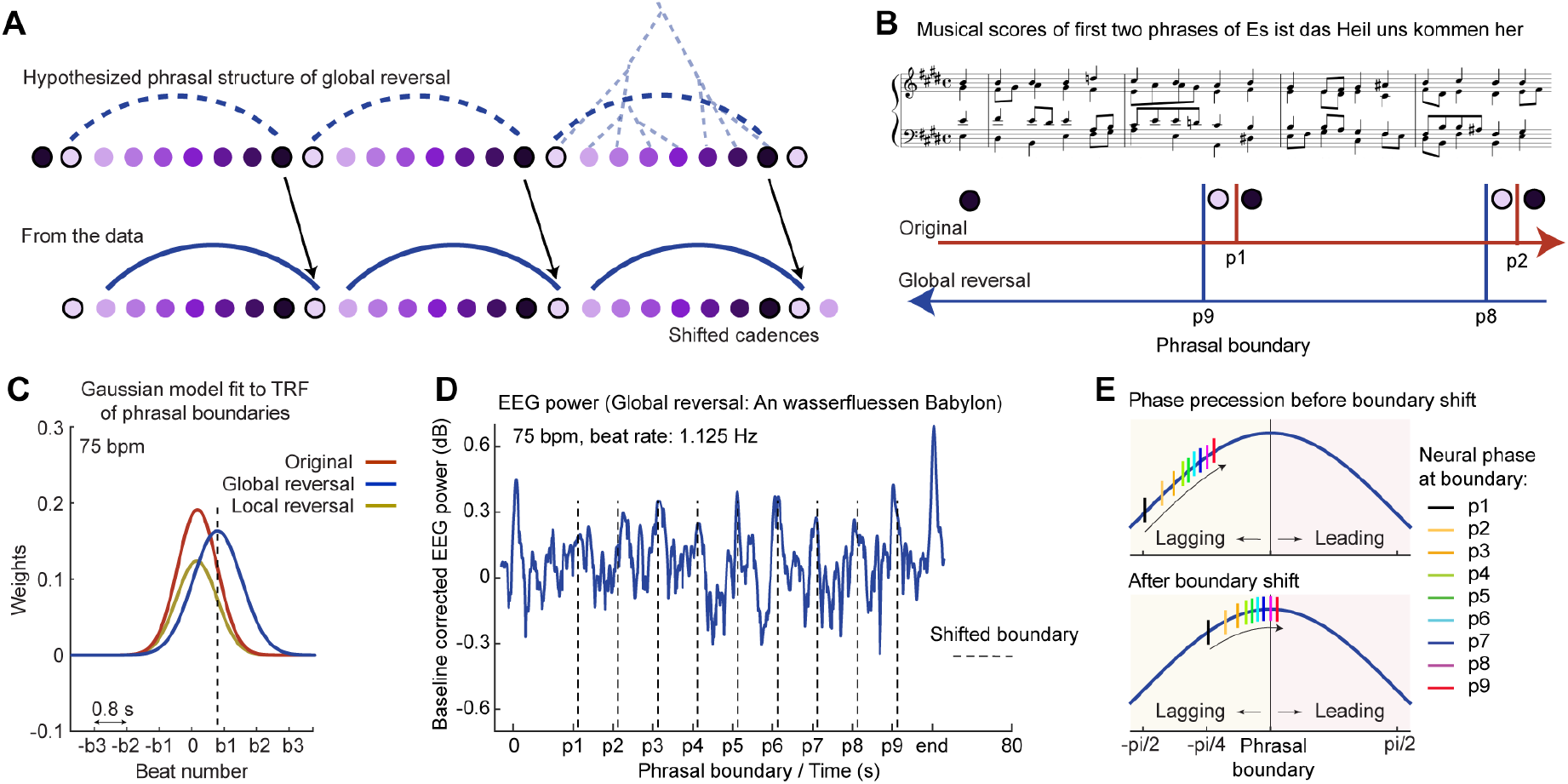
Phrasal segmentation in the Global reversal condition, related to Fig. 3&4. (A) Illustration of phrasal segmentation in the Global reversal condition. We hypothesized in the beginning that the phrasal structures in the Global reversal condition preserved the original phrasal boundaries (upper part of the plot). However, the data in Fig. 3&S4 suggest that the neural signals did not lock to the original phrasal boundaries but shifted forward, which was probably due to the fact that the brain segments a musical phrase after the end beat of a musical phrase in the Original condition, which was the onset beat in the Global reversal condition according to the pre-defined phrasal boundaries (lower part of the plot). This means that the phrasal boundaries in the Global reversal condition shifted one beat forward. See Table S2 for the cadences of the Global reversal condition. (B) Illustration of phasal segmentation in the Global reversal through an example excerpt. In the Original condition, phrasal boundaries end after the beat marked by purple-filled circle (potentially a strong beat); in the Global reversal condition, phrasal boundaries end after the beat marked by purple-filled circle. But, as the beat order was reversed in the Global reversal, the phrasal boundaries shifted one beat forward. (C) TRFs of phrasal boundaries at 75 bpm. The peak latency of TRF in the Global reversal was close to the beat position of ‘b1’, instead of around ‘0’ - the pre-defined phrasal boundary in the Original condition and the Local reversal condition. This indeed suggests that the phrasal boundaries shifted one beat forward in the Global reversal condition. (D) EEG power of an example music piece in the Global reversal condition. The x axis marks the pre-defined phrasal boundaries. The dashed line represents the new phrasal boundaries shifted forward by one beat. Indeed, the data show that the neural signals were advancing as music unfolded and led over the new phrasal boundaries by the end of the music piece. (E) Illustration of neural phase shift in the Global reversal condition. We plot the neural phases similarly as in Fig. 5B, but in the upper panel using the pre-defined phrasal boundaries whereas in the lower panel using the new phrasal boundaries. It can be clearly seen in the lower panel that the neural phrases led over the phrasal boundaries by the end of the music piece.

**Fig. S6.**
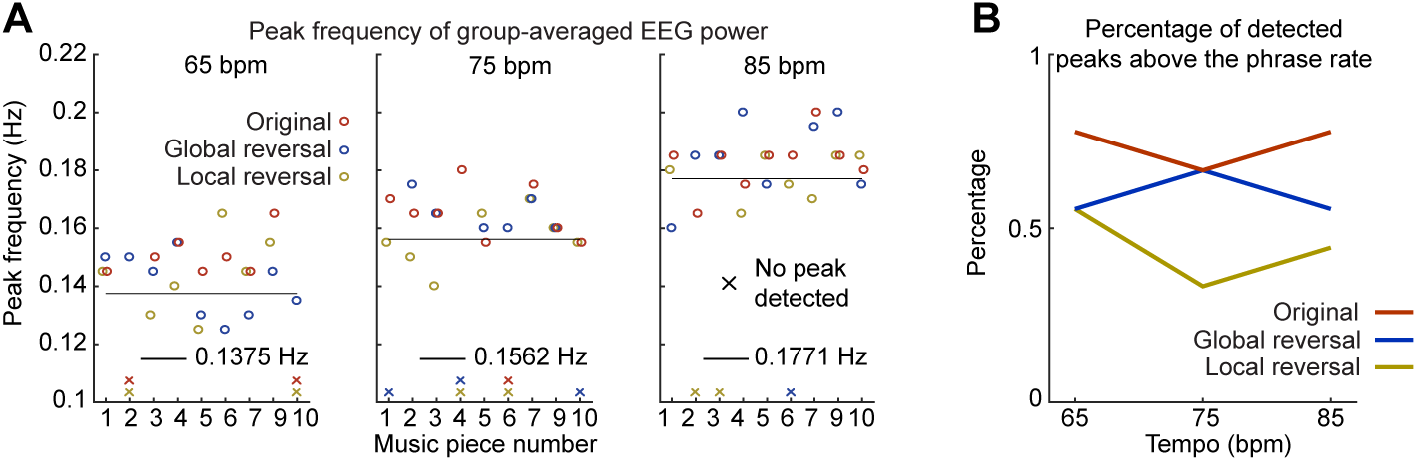
Peak frequency of group-averaged EEG power, related to Fig. 4. (A) We automatically detected the peak frequency of the phrasal segmentation for each music piece within the significant frequency ranges around F0 defined in Fig. 3C. The circle color codes for the reversal conditions. The crosses right above the x axis indicate the music pieces in which no peak frequency was able to be detected using our algorithm. The thin black line indicates the phrase rate at each tempo. (B) We calculated the percentage of the peak frequencies that are above the phrase rate at each tempo for each condition. The y axis represents the percentage with ‘1’ indicating that all the peak frequencies are above the phrase rate. The data is consistent with the results of PPPi in Fig. 5, showing that, when the phrase precession occurred, the peak frequency of the phrasal segmentation was larger than the phrase rate of the music pieces.

**Fig. S7.**
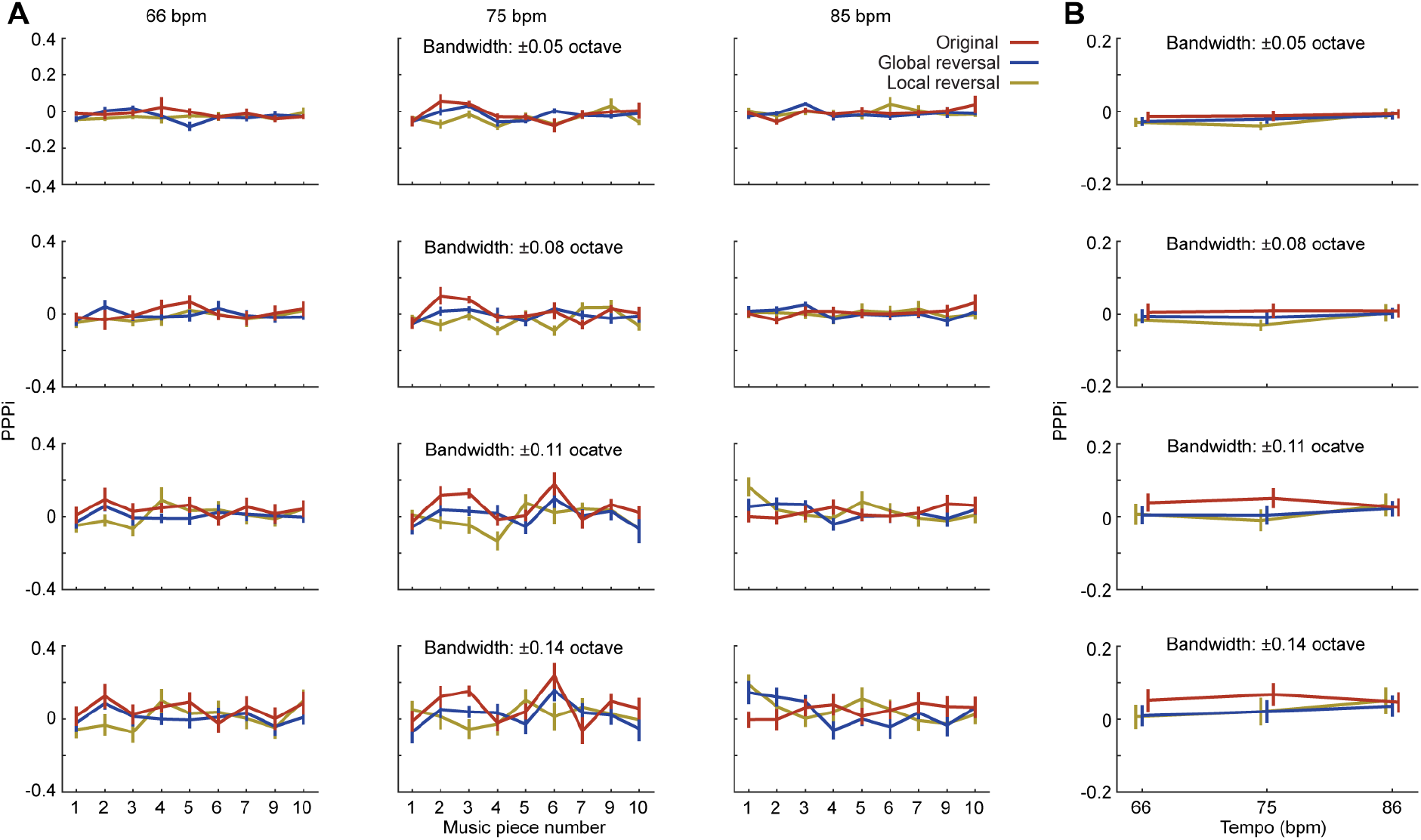
PPPi derived from individual participants at different bandwidths (see Methods), related to Fig. 4. (A) PPPi of each music piece from individual participants. We followed the same analysis procedure as in Fig. 5F but calculated PPPi for each participant first and then averaged the PPPi values over all the participants. The bandwidth used to calculate PPPi was varied systemically from ± 0.05 octave around the phrase rate at each tempo to ± 0.14 octave. The error bars represent ±1 standard error of mean over the participants. (B) Averaged PPPi for each condition. We averaged PPPi over nine music pieces in each condition for each participant and then conducted a two-way rmANOVA (Tempo * Condition) at each bandwidth over participants. The error bars represent ±1 standard error of mean over the participants. 1) Bandwidth ± 0.05 octave (65 bpm: 0.1328 – 0.1423 Hz; 75 bpm: 0.1509 – 0.1617 Hz; 85 bpm: 0.1711 – 0.1833 Hz): Tempo (*F*(2,56) = 7.27, *p* = 0.002, η_p_^2^ = 0.206); Condition (*F*(2,56) = 2.72, *p* = 0.075, η_p_^2^ = 0.089); the interaction (*F*(4,112) = 1.20, *p* = 0.314, η_p_^2^ = 0.041). 2) Bandwidth ± 0.08 octave (65 bpm: 0.1301 – 0.1453 Hz; 75 bpm: 0.1478 – 0.1651 Hz; 85 bpm: 0.1675 – 0.1872 Hz): Tempo (*F*(2,56) = 1.94, *p* = 0.153, η_p_^2^ = 0.065); Condition (*F*(2,56) = 3.75, *p* = 0.030, η_p_^2^ = 0.118); the interaction (*F*(4,112) = 0.60, *p* = 0.664, η_p_^2^ = 0.021). 3) Bandwidth ± 0.11 octave (65 bpm: 0.1274 – 0.1484 Hz; 75 bpm: 0.1447 – 0.1686 Hz; 85 bpm: 0.1641 – 0.1911 Hz): Tempo (*F*(2,56) = 0.78, *p* = 0.463, η_p_^2^ = 0.027); Condition (*F*(2,56) = 3.63, *p* = 0.033, η_p_^2^ = 0.115); the interaction (*F*(4,112) = 1.26, *p* = 0.291, η_p_^2^ = 0.043). 4) Bandwidth ± 0.14 octave (65 bpm: 0.1248 – 0.1515 Hz; 75 bpm: 0.1418 – 0.1721 Hz; 85 bpm: 0.1604 – 0.1951 Hz): Tempo (*F*(2,56) = 1.32, *p* = 0.276, η_p_^2^ = 0.045); Condition (*F*(2,56) = 3.14, *p* = 0.051, η_p_^2^ = 0.101); the interaction (*F*(4,112) = 0.65, *p* = 0.628, η_p_^2^ = 0.023). When the bandwidth is narrow (± 0.05 octave), the main effect of Tempo is significant. When the bandwidth increased, the main effect of Tempo was not significant anymore. This potentially explained that the effect of the Tempo on PPPi was modulated or driven by the bandwidth. As the spectral range of the phrasal segmentation varied with the tempi, the frequency range for calculating PPPi needs to properly cover the spectral components relevant to the phrasal segmentation. When the bandwidth was too narrow, it distorted the estimation of PPPi – the center frequency of a frequency band used at the large tempo fell below the peak frequencies of the music pieces, so the peak frequencies of all the conditions are seemingly larger than the center frequency and the PPPi was overestimated. The main effect of Condition is significant when the bandwidth is ± 0.08 octave (*p* = 0.030) or ± 0.11 octave (*p* = 0.033), but not when the bandwidth is ± 0.05 octave (*p* = 0.075) or ± 0.014 octave (*p* = 0.051). The reason is probably that, if the bandwidth is too narrow, the spectral components relevant to the phrasal segmentation were not fully included; if the bandwidth is too wide, the frequency range included the spectral components that were unrelated to the phrasal segmentation. Note that, in Fig. 5, the bandwidth was not determined arbitrarily or varied manually but was derived in a data-driven way from the significant frequency ranges in Fig. 4C.

**Table S1.** Selected chorales and their predefined phrasal structures marked by fermatas.

| No. | Piece code | Piece Name | Phasal structure (beats per phrase) |
| --- | --- | --- | --- |
| 0 | BWV 255 | Ach Gott und Herr | 4 4 8 4 4 8 |
| 1 | BWV153/1 | Ach Gott, vom Himmel sieh darein | 8 8 8 8 8 8 8 |
| 2 | BWV 3/6 | Ach Gott, wie manches Herzeleid | 8 8 8 8 |
| 3 | BWV267 | An Wasserflüssen Babylon | 8 8 8 8 8 8 8 8 12 |
| 4 | BWV 6/6 | Erhalt uns Herr, bei deinem Wort | 8 8 8 8 |
| 5 | BWV 248 | Ermuntre dich, mein schwacher Geist | 8 8 8 8 8 8 8 8 |
| 6 | BWV 86/6 | Es ist das Heil uns kommen her | 8 8 8 8 8 8 8 |
| 7 | BWV 308 | Es spricht der Unweisen Mund wohl | 8 8 8 8 8 8 8 |
| 8 | BWV 20/7 11 | O Ewigkeit, du Donnerwort | 12 12 12 12 12 12 |
| 9 | BWV 245/5 | Vater unser im Himmelreich | 8 8 8 8 8 8 8 8 |
| 10 | BWV 110/7 | Wir Christenleut | 8 8 8 8 8 8 |

**Table S2.** Musical segment boundaries determined according to voice leading, cadence, and group preference rule. The numbers below refer to the number of beats within a segment determined by each musical criterion.

| Piece number and condition | Voice leading | Cadence | Grouping rule |
| --- | --- | --- | --- |
| 1 Original | 4 4 4 4 4 4 4 8 8 8 | 4 4 4 4 4 4 4 4 4 4 4 8 | 4 4 4 4 4 4 4 4 4 4 4 5 7 |
| 1 Global reversal | 4 4 6 2 6 2 5 4 4 3 5 4 4 3 | 8 5 9 7 4 4 3 5 4 4 3 | 8 8 6 3 4 3 5 4 4 3 5 3 |
| 1 Local reversal | 5 3 5 3 5 3 5 3 6 2 6 2 4 4 | 5 3 5 3 5 3 5 3 6 2 6 2 8 | 5 3 5 3 5 3 5 3 6 2 6 2 8 |
| 2 Original | 8 8 8 8 | 8 8 8 8 | 3 5 8 8 8 |
| 2 Global reversal | 8 8 8 8 | 8 8 9 7 | 2 6 8 2 7 7 |
| 2 Local reversal | 8 8 8 8 | 8 8 8 8 | 8 8 8 8 |
| 3 Original | 8 8 8 8 8 8 8 8 12 | 8 8 8 8 8 8 8 8 12 | 8 8 8 8 8 8 8 8 12 |
| 3 Global reversal | 12 9 9 15 7 9 7 9 7 | 12 9 9 8 7 7 9 7 9 7 | 12 8 8 8 8 8 8 8 8 |
| 3 Local reversal | 8 8 8 8 8 7 9 8 7 13 | 8 8 8 8 4 4 12 4 8 8 8 4 | 8 8 8 8 8 8 4 4 8 8 12 |
| 4 Original | 8 8 8 8 | 8 8 8 8 | 8 8 8 8 |
| 4 Global reversal | 7 9 4 4 8 | 7 5 4 6 4 6 | 2 6 4 4 4 4 4 4 |
| 4 Local reversal | 8 6 10 8 | 8 9 7 8 | 8 8 8 8 |
| 5 Original | 8 8 8 8 8 8 8 8 | 8 8 8 8 8 8 8 8 | 8 5 3 8 5 3 5 3 8 8 8 |
| 5 Global reversal | 8 11 6 8 10 6 10 5 | 8 11 8 6 10 6 10 5 | 8 8 8 8 8 8 8 8 |
| 5 Local reversal | 7 8 8 8 8 9 8 | 16 16 7 8 10 7 | 2 5 3 6 2 5 3 6 8 2 6 8 4 4 |
| 6 Original | 4 4 4 4 4 4 4 4 8 8 8 | 4 4 8 4 4 8 8 8 8 | 4 4 8 4 4 8 9 7 8 |
| 6 Global reversal | 8 8 8 6 10 6 10 | 8 8 5 3 4 4 4 4 4 4 4 4 | 3 5 3 5 4 4 4 4 4 4 4 4 4 4 |
| 6 Local reversal | 7 7 9 7 10 6 10 | 5 3 5 3 5 3 5 3 6 8 6 4 | 4 4 4 4 4 4 4 4 4 4 4 4 4 4 |
| 7 Original | 8 8 8 8 8 8 8 | 8 4 4 8 4 4 8 8 8 | 4 4 4 5 3 4 4 5 3 4 8 3 5 |
| 7 Global reversal | 8 7 8 8 9 7 9 | 9 8 6 8 5 4 7 5 4 | 4 4 8 4 4 3 4 5 4 3 4 5 4 |
| 7 Local reversal | 8 7 9 7 9 7 9 | 8 8 8 8 8 8 8 | 8 9 7 9 7 8 8 |
| 8 Original | 12 12 12 12 12 12 | 12 12 4 5 3 12 12 12 | 9 3 12 4 8 12 12 12 |
| 8 Global reversal | 8 11 6 11 8 8 8 12 | 12 13 11 13 11 12 | 12 12 12 12 12 12 |
| 8 Local reversal | 4 4 4 8 4 8 5 8 4 11 8 4 | 12 12 12 13 11 12 | 3 9 12 8 4 12 12 12 |
| 9 Original | 8 8 8 8 8 8 8 8 | 8 8 8 8 8 8 8 8 | 8 8 8 8 8 8 8 8 |
| 9 Global reversal | 4 4 8 8 4 4 8 8 4 4 8 | 8 8 8 8 8 8 8 8 | 4 4 8 8 8 8 8 8 8 |
| 9 Local reversal | 1 7 8 8 1 7 8 8 8 4 4 | 8 8 8 8 8 8 8 8 | 4 4 8 4 4 4 4 8 4 4 4 4 8 |
| 10 Original | 8 8 8 8 8 8 | 8 8 8 8 8 8 | 8 8 8 8 8 8 |
| 10 Global reversal | 8 7 12 8 8 5 | 8 7 3 3 5 17 5 | 3 5 4 4 8 4 4 4 4 8 |
| 10 Local reversal | 8 8 9 7 7 9 | 8 8 9 7 7 9 | 8 8 8 8 8 8 |

**Table S3.** Correlations between the neural tracking at each tempo and the score at the music training subscale of the GOLD-MSI. As the main effect of Condition was not significant (see main text), we averaged Cacoh values over the conditions before computing the correlation.

| Beat rate |  |  |  |
| --- | --- | --- | --- |
| Tempo (bpm) | 65 | 75 | 85 |
| r | 0.366 | 0.327 | 0.317 |
| p value | 0.050 | 0.082 | 0.093 |
| Note rate |  |  |  |
| Tempo (bpm) | 65 | 75 | 85 |
| r | 0.465 | 0.391 | 0.439 |
| p value | 0.010 | 0.035 | 0.017 |

**Table S4.** Correlations between RMS in the early and late regions and the score at the music training subscale of the GOLD-MSI for each tempo and each condition.

| Early period (0.03 - 0.1 s) |  |  |  |  |  |  |  |  |  |
| --- | --- | --- | --- | --- | --- | --- | --- | --- | --- |
| Tempo (bpm) | 65 |  |  | 75 |  |  | 85 |  |  |
| Condition | Local | Global | Original | Local | Global | Original | Local | Global | Original |
| r | 0.349 | 0.351 | 0.445 | 0.428 | 0.376 | 0.316 | 0.328 | 0.402 | 0.372 |
| p value | 0.063 | 0.062 | 0.016 | 0.021 | 0.045 | 0.095 | 0.082 | 0.031 | 0.047 |

| Late period (0.12 - 0.25 s) |  |  |  |  |  |  |  |  |  |
| --- | --- | --- | --- | --- | --- | --- | --- | --- | --- |
| Tempo (bpm) | 65 |  |  | 75 |  |  | 85 |  |  |
| Condition | Local | Global | Original | Local | Global | Original | Local | Global | Original |
| r | 0.255 | 0.276 | 0.361 | 0.314 | 0.304 | 0.251 | 0.279 | 0.308 | 0.278 |
| p value | 0.182 | 0.148 | 0.055 | 0.098 | 0.109 | 0.188 | 0.142 | 0.104 | 0.145 |

